# Heterogeneous migration of neuronal progenitors to the insula shapes the human brain

**DOI:** 10.1101/2022.09.09.507371

**Authors:** Arka N. Mallela, Hansen Deng, Ali Gholipour, Simon K Warfield, Ezequiel Goldschmidt

## Abstract

The human cerebrum consists of a precise and stereotyped arrangement of lobes, gyri, and connectivity that underlies human cognition. The development of this arrangement is less clear. Current models of radial glial cell migration explain individual gyral formation but fail to explain the global configuration of the cerebral lobes. Moreover, the insula, buried in the depths of the Sylvian fissure, belies conventional models. Here, we show that the insula has unique morphology in adults, that insular morphology and slow volumetric growth emerge during fetal development, and that a novel theory of curved migration is required to explain these findings. We calculated morphologic data in the insula and other lobes in adults (N=107) and in an *in utero* fetal brain atlas (N=81 healthy fetuses). *In utero*, the insula grows an order of magnitude slower than the other lobes and demonstrates shallower sulci, less curvature, and less surface complexity both in adults and progressively throughout fetal development. Novel spherical projection analysis demonstrates that the lenticular nuclei obstruct 60-70% of radial pathways from the ventricular zone (VZ) to the insula, forcing a curved migration path to the insula in contrast to a direct radial pathway. Using fetal diffusion tractography, we identify streams of putative progenitor cells that originate from the VZ and migrate tangentially *around* the lenticular nuclei to form the insula. These results challenge existing models of radial migration to the cortex, provide an alternative model for insular and cerebral development, and lay the groundwork to understand cerebral malformations, insular functional connectivity, and insular pathologies.

## INTRODUCTION

The complex folding of the human cerebral cortex underlies its capacity for remarkable information processing, cognitive and sensorimotor functions. The characteristic gyral patterns observed in the cortex have been attributed to tangential expansion and long-distance radial migration along radial glial scaffolds.^1–3^ Neural progenitors located in transient embryonic zones, including the ventricular zone (VZ) and subventricular zone (SVZ), proliferate and move across the intermediate zone through this scaffolding.^4–7^ More recent observations suggest a possible two-step model of neurogenesis, first with radial glia undergoing asymmetric division in the VZ, followed by intermediate progenitor cells in the SVZ producing neurons through symmetric division.^8–11^ The process of radial migration was previously thought to be complete by GA 24 weeks, but recent evidence suggests that this process continues after birth into infancy.^12^

While gyral formation through radial growth does characterize local folding, it does not explain the development of the global structure of the brain and the arrangement of the cerebral lobes. Landmark studies that have explored the mechanical process of gyrification have largely assumed a homogenous pattern growth of the brain.^13–15^ Simulations *in silico* and in gel models suggest that constrained cortical expansion can lead to folding patterns that are similar to human sulci and gyri.^14,16^ However, these models largely do not account for the formation of the insula. Buried in the depths of the Sylvian fissure, the insula is involved in interoceptive function and interfaces with cortical and subcortical sensorimotor and limbic areas.^17^ While the insula does have a stereotyped pattern of short and long gyri, insular gyri are shallower, straighter, and less complex compared to those of the cerebral convexity ^18^.

The insula is also the focal point around which the Sylvian fissure, and accordingly the operculae of the frontal, parietal, and temporal lobes, are arranged.^18–20^ Here we show that the distinct C-shape of the cerebrum is a function of these operculae closing over the insula during fetal development. Accordingly, understanding the determinants of how the other lobes overgrow the insula to close to Sylvian fissure is paramount to understanding the global configuration of cerebrum.^20^ Current models of radial growth fail to explain both the unique morphology and position of the insula and the arrangement of the cerebral lobes around the Sylvian fissure.^21,22^

Previously, we have shown that the Sylvian fissure forms through a unique process of folding compared with other sulci^20^ and that these differences may be due to distinct transcriptional signatures in the insula vs. the opercula.^23^ Given the anatomic and morphological differences of the insula, we hypothesized that the development of the insula does not follow the model of radial migration that forms the other lobes, and that the differences in the mechanisms driving insular and opercular growth determine the overall shape of the human cerebrum. In this study, we quantify the morphologic differences between the insula and the other lobes in adults and identify that these differences emerge during fetal development. We then show that existing models of radial migration cannot explain the formation of the insula. In contrast, we identify a distinct curved and tangential migratory route of developing cells to the insular cortex, in contrast to radial pathways are found in all other lobes. This heterogeneity in radial migration yields different lobe volumes, folding pattern, and ultimately shapes the surface of the human brain.

## RESULTS

### Adult insula anatomy

Our investigation stemmed from the observation that the adult insula is anatomically and morphologically different than the other lobes of the brain.^18,19^ The insula is covered by the frontal, temporal, and parietal operculae in the Sylvian fissure and is not apparent on the lateral cerebral surface of the adult brain **(Fig. 1a).** Furthermore, upon exposure, the sulci (and gyri) of the insula immediately appear to be shallower and less convoluted than those on the other lobes **(Fig. 1b).** We sought to quantify these differences. Using N=107 adult patients (54.2% female) of median age 28 (range 22-35) from the Human Connectome Project HCP 900 subjects data release^24^, we quantified surface area **(Fig. 1d),** range from deepest sulcal fundus to most superficial gyral crown (sulcal depth range [SDR]) **(Fig. 1e),** extrinsic (mean) curvature **(Fig. 1f),** fractal dimension (FD) – a measure of surface complexity^25–27^ **(Fig. 1g)**, cortical thickness **(Supplementary Fig. 2d),** and intrinsic (Gaussian) curvature **(Supplementary Fig. 2f).** We found significant differences between the insula and other lobes (frontal, parietal, temporal and occipital) in every parameter. In particular, the insula had smaller surface area (surface area=26.7 vs. frontal 333.9/parietal 243.8/temporal 186.2/occipital 115.7 cm^2^, p<0.0001 pairwise against all lobes), shallower sulci (SDR=2.53 vs. 3.23/3.34/3.32/2.95 cm, p<0.0001), had less extrinsic curvature – (curvature=0.0 vs. 0.05/0.05/0.05/0.06 mm^-1^, p<0.0001), and less complex surface (Fractal Dimension=2.25 vs. 2.48/2.46/2.49/2.40, p<0.0001), than the other lobe. These findings were in concordance with the anatomic studies of the insula. To better understand this high dimensional dataset, we utilized UMAP (Uniform Manifold Approximation and Projection) to visualize these data, clearly demonstrating a separate clustering of the insula **(Fig. 1c).** As the insula is smaller in adults, surface area was not included in the UMAP to avoid biasing results.

**Fig. 1:**
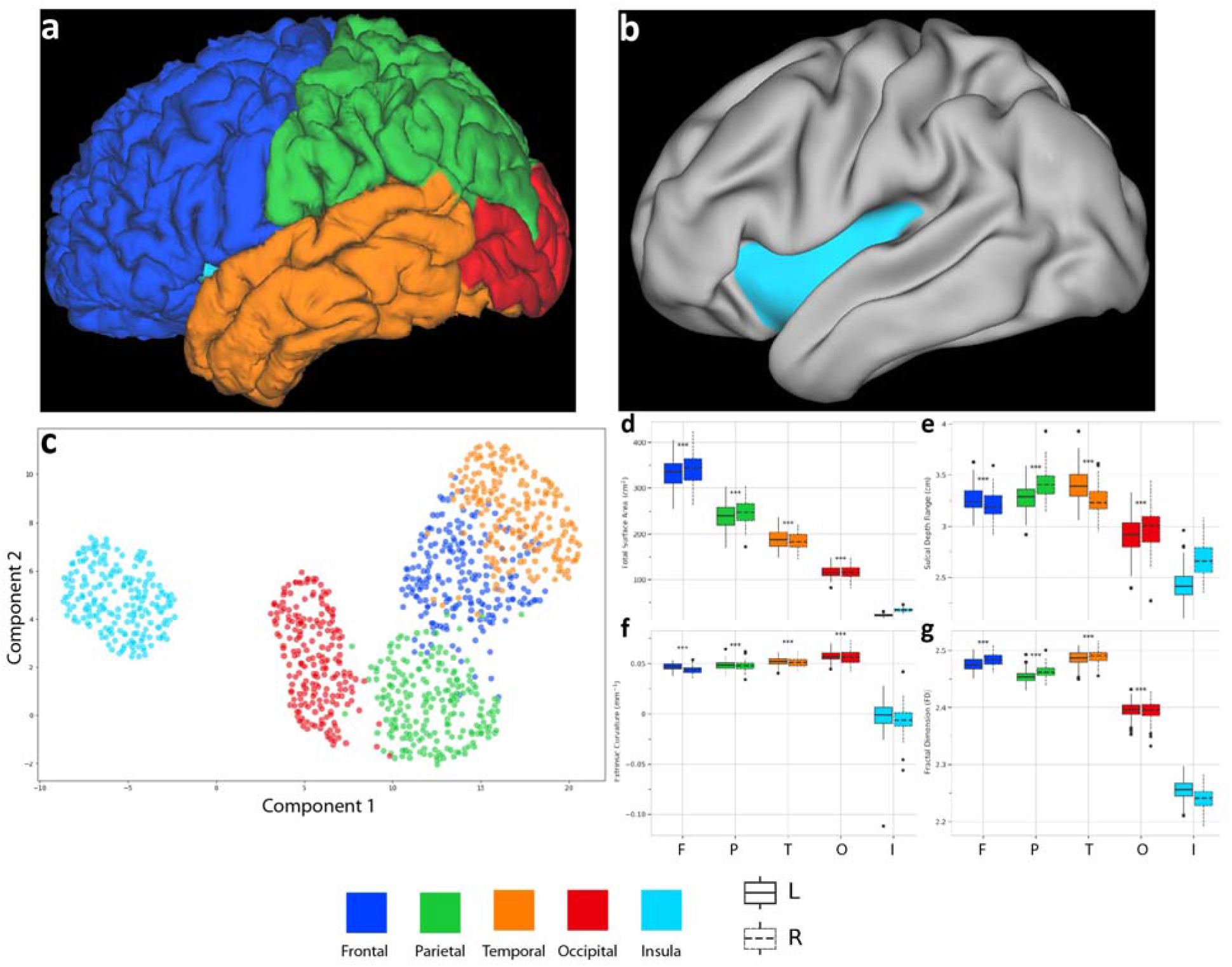
The insula is morphologically different in adults. **a:** Adult pial surface. The convoluted surface of the frontal (blue), parietal (green), temporal (orange), and occipital (red) lobes stands in stark contrast to that of the insula. In unexpanded view, the insula is entirely covered by the frontal, parietal, and temporal operculae. **b:** Anatomic specimen demonstrating insula and surrounding operculae. After retractors are placed, the convoluted gyration of the frontal and temporal contrast to the relatively shallow and straight gyri of the insula. White arrow – central sulcus of insula. **c:** UMAP of cortical surface metrics (cortical thickness, sulcal depth range, mean curvature, fractal dimension) in N=107 adult subjects. The insula clusters separately. UMAP parameters: # of neighbors = 20, minimum distance = 0.7, seed = 281. **d:** Total surface area by lobe. Linear regression against side and lobe was significant with high overall fit (R^2^ = 0.983, F(6,1064)=16460, p<0.0001). Median surface area in the frontal (339.9 cm^2^; p < 0.0001), parietal (243.8 cm^2^; p < 0.0001), temporal (186.2 cm^2^; p < 0.0001), and occipital (115.7 cm^2^; p < 0.0001) cortices was significantly higher than the insula (26.7 cm^2^), using pair-wise Mann-Whitney testing with Bonferroni correction. **e:** Sulcal depth range (SDR) - range from deepest sulcal fundus to most superficial gyral crown, by lobe. Linear regression against side and lobe was significant with high overall fit (R^2^ = 0.997, F(6,1064)=61070, p<0.0001). Median SDR in the frontal (3.23 cm; p < 0.0001), parietal (3.34 cm; p < 0.0001), temporal (3.32 cm; p < 0.0001), and occipital (2.95 cm; p < 0.0001) cortices was significantly higher than the insula (2.53 cm). **f:** Extrinsic (mean) curvature by lobe. Linear regression against side and lobe was significant with high overall fit (R^2^ = 0.973, F(6,1064)=6501, p<0.0001). Median curvature in the frontal (0.05 mm^-1^; p < 0.0001), parietal (0.05 mm^-1^; p < 0.0001), temporal (0.05 mm^-1^; p < 0.0001), and occipital (0.06 mm^-1^; p < 0.0001) cortices was significantly higher than the insula (0.0 mm^-1^). **g:** Fractal dimension (FD) by lobe. Linear regression against side and lobe was significant with high overall fit (R^2^ = 1.00, F(6,1064)=503400, p<0.0001). Median FD in the frontal (2.48; p < 0.0001), parietal (2.46; p < 0.0001), temporal (2.49; p < 0.0001), and occipital (2.40; p < 0.0001) cortices was significantly higher than the insula (2.25). *** - p < 0.0001 (Mann-Whitney Test – lobe vs. Insula, Bonferroni corrected)

### Fetal insular development

Given the differences in adult anatomy, we hypothesized that the unique morphology of the insula emerged *in utero*. To investigate this, we utilized the CRL fetal brain MRI atlas developed by Gholipour et al.^28^ constructed from weekly fetal T2-weighted *in utero* MRI images of 81 healthy fetuses between 19 and 39 weeks of gestational age (GA). Unlike in the adult subjects, the insula is readily apparent on the lateral cortical surface during mid second trimester and progressively is covered by the frontal, parietal, and temporal opercula during the third trimester **(Supplementary Fig. 1, Fig. 2a).** Moreover, visually, the insula remains relatively free of sulci and gyri until the appearance of the central sulcus of the insula until late in gestation, approximately at GA 36 weeks **(Fig. 2b).** This lent credence to our hypothesis that the differences in insula morphology emerged during gestation.

**Fig. 2:**
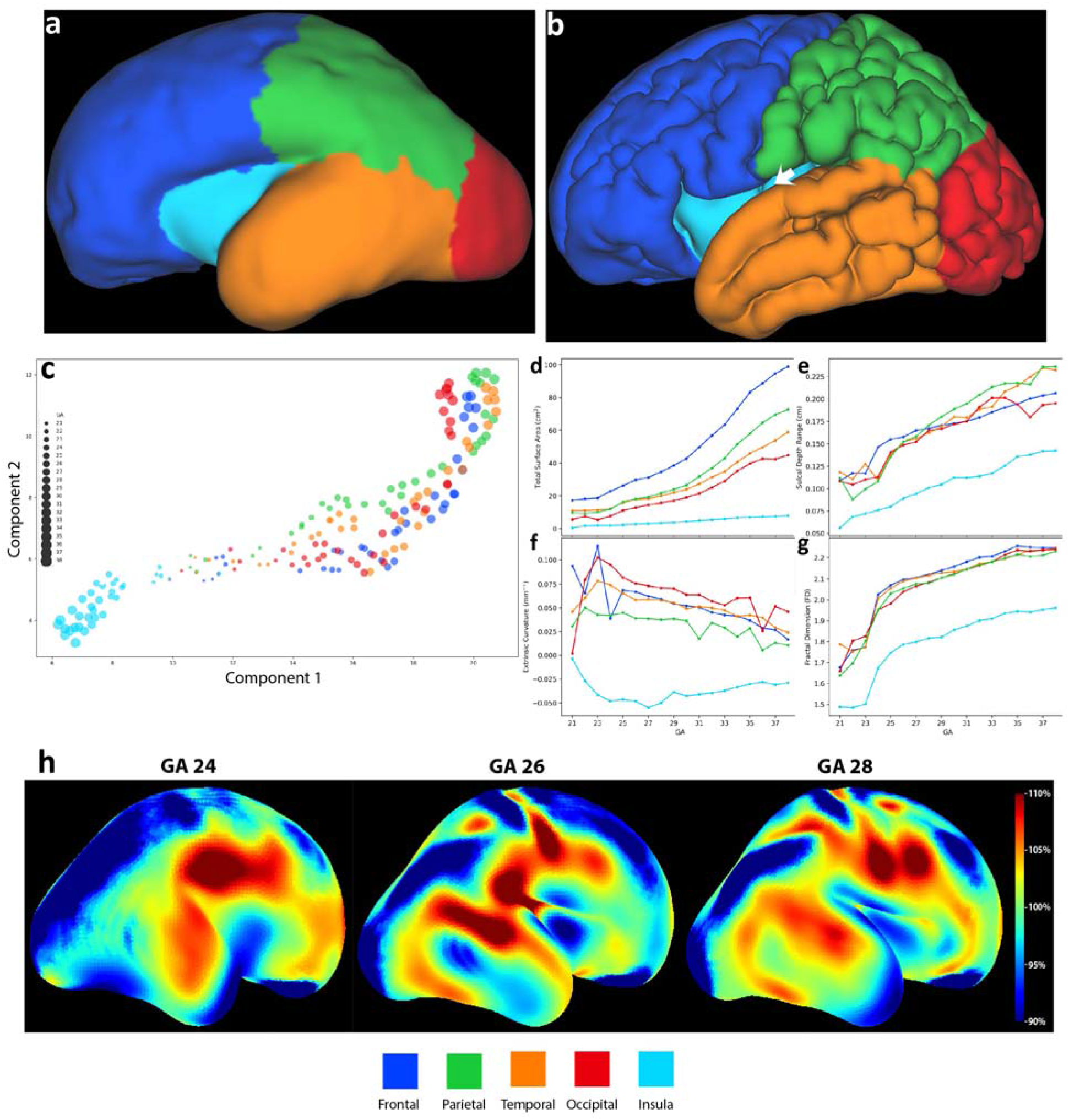
Altered insular morphology and growth patterns emerge during fetal development and drive formation of the Sylvian fissure and cerebral convexity. **a:** Pial surface of the left hemisphere at GA 24 weeks from the CRL fetal brain MRI atlas. The entire brain lacks gyri, and the insula is uncovered. **b**: Pial surface of the left hemisphere at GA 36 weeks. The frontal, temporal, parietal, and occipital lobes have developed significant gyri and sulci and the operculae have begun to cover the insula. In contrast the insula remains largely flat and devoid of significant sulci, except for the shallow central sulcus of the insula (white arrow), which appears at GA 35-36 weeks. **c**: UMAP of cortical surface metrics (cortical thickness, sulcal depth range, mean curvature, and fractal dimension) across fetal development. Larger circles indicate later stages in development. Lobe volume and surface area are excluded. The insula takes a different trajectory than the other lobes. UMAP parameters: # of neighbors = 75, minimum distance = 0.4, seed = 74. **d:** Surface area by lobe by GA. Surface area grows exponentially for frontal, temporal, parietal, and occipital lobes, but linearly in the insula (Supplementary Table 2). Linear regression against side, GA, and lobe was significant with good overall fit (R^2^ = 0.841, F(6,173)=159.0, p<0.0001). Surface area was significantly higher in the frontal, parietal, temporal, and occipital cortices than the insula (p < 0.001). **e:** Sulcal depth range by GA. Linear regression against side, GA, and lobe was significant with good overall fit (R^2^ = 0.927, F(6,173)=377.5, p<0.0001). SDR was significantly higher in the frontal, parietal, temporal, and occipital cortices than the insula (p < 0.001). **f:** Extrinsic (mean) curvature by GA. Linear regression against side, GA, and lobe was significant with moderate fit (R^2^ = 0.487, F(6,173)=29.33, p<0.0001). Curvature was significantly higher in the frontal, parietal, temporal, and occipital cortices than the insula (p < 0.001). **g:** Fractal dimension by GA. Linear regression against side, GA, and lobe was significant with good overall fit (R^2^ = 0.852, F(6,173)=173.3, p<0.0001). FD was significantly higher in the frontal, parietal, temporal, and occipital cortices than the insula (p < 0.001). **h:** Jacobian analysis (week to week % relative volume expansion, correcting for overall volume growth) of the cerebrum (right lateral view at GA 24, 26, 28) demonstrates significantly increased growth in the frontal, temporal, and parietal operculae compared to the insula. Over the second and third trimester, this leads to the closure of the Sylvian fissure over the insula and creates the characteristic C-shape of the cerebrum^20^.

The morphological differences observed in adults were also found in developing fetal brain. Specifically, while the insula is similar in surface area to the other lobes at GA 21 weeks, it is significantly smaller than the other lobes by birth at GA 38 weeks (p<0.0001), concordant with adults **(Fig. 2d).** We used curve fitting to model the type of growth (logistic, exponential, linear) in surface area and other parameters. The surface area in all lobes grew exponentially except for the insula which grew linearly **(Supplementary Table 2).** Sulcal depth range **(Fig. 2e),** extrinsic curvature **(Fig. 2f),** and fractal dimension **(Fig. 2g)** were all lower in the insula vs. the other lobes at GA 21 weeks and remained so during gestation until birth (p<0.0001, linear regression against side, GA, and lobe). Interestingly, fractal dimension demonstrated an inflection point in all lobes at approximately GA 24 weeks but remained smaller in the insula. Unlike in the adult cohort, intrinsic curvature and cortical thickness did not show significant differences by lobe during fetal development **(Supplementary Fig. 2c,** f).

To visualize the high dimensional dataset of fetal morphometric data, we again utilized UMAP (excluding volume and surface area) to visualize the results by lobe **(Fig. 2c).** The insula clearly clustered differently than the other lobes. Moreover, the insula cluster diverges from the other lobes by gestational age. These analyses indicate that the morphological differences observed in the insula in adults emerge during fetal development.

We also performed volumetric analysis by lobe during fetal development. Each lobe is associated with white matter that constitutes the bulk of the volume of that lobe. In contrast, the insula lacks a deep white matter and only has superficial white matter in the extreme capsule. In concordance with anatomic studies^19^, we treated the central core of the brain (defined as the caudate nucleus, lenticular nuclei – putamen and globus pallidus, internal capsule) as the deep structures of the insula. The volume of the frontal lobe was 5.0 cc at 21 weeks and grew 8.5-fold to 42.7 cc at 38 weeks. Similarly, the temporal lobe grew 7-fold, the parietal lobe 7.5-fold, and the occipital lobe 7-fold. In contrast, the insula and central core grew from 1.8 cc at 21 weeks to only 9.2 cc at Week 38, a 5.1-fold absolute increase **(Supplementary Fig. 2a).** Growth modeling revealed that all lobes exhibited logistic growth. However, the logistic growth rate *α* was an order of magnitude lower in the insula and central core than the other lobes (0.002 vs. 0.05/0.03/0.04/0.02; **Supplementary Table 2).** Growth in the central core was driven by deep gray matter - the caudate (0.6 to 1.5cc) and lenticular nuclei (0.5 to 2.5cc), white matter (0.3 to 1.9cc), and the insular cortex itself (0.4 to 3.3cc). **(Supplementary Fig. 2b).**

We then performed Jacobian volumetric analysis, correcting for overall cerebral growth, to localize the regions within each lobe that grew most and the overall changes in the shape of the cerebrum through development. Specifically, from approximately weeks 23-28, frontal, temporal, and parietal operculae demonstrate 110% volumetric growth per week compared to overall growth, while the insula relatively shrinks (90% compared to overall cerebral growth). This process progressively expands the frontal, parietal, and temporal lobes over the insula, closing the Sylvian fissure **(Fig. 2g).** The relative slow growth in the insula (in volume and surface area) is responsible for this process and accordingly the overall configuration of the human cerebrum.

### Origin of insular progenitors

We subsequently investigated the development origins of this stark difference in insular morphology and growth pattern. The cerebral cortex emerges from radially migrating neural progenitors and radial expansion is at the center of sulcal morphogenesis.^12,29^ Progenitor cells travel from the ventricular zone (VZ) along radial glial cells by way of the subventricular zone (SVZ).^29^ In the majority of the cerebrum, there is a direct path from the VZ/SVZ through developing white matter to the cortex. In contrast, there are several key deep gray matter structures that could obstruct direct radial migration of neural progenitor cells along radial glia from the VZ to the insula, particularly the developing globus pallidus and putamen (collectively lenticular nuclei) **(Supplementary Fig. 1**).

We hypothesized that progenitor cells for each cortex traversed a linear path from the closest point in the ventricular zone but that the path to the insula is blocked by the lenticular nuclei, preventing a radial trajectory. We segmented the VZ by the closest lobe (in Euclidean distance) **(Supplementary Fig. 3a).** The segmentation largely follows the expected anatomic course, with the VZ surrounding the frontal horn and body of the lateral ventricle corresponding to the frontal lobe, atrium to the parietal, occipital horn to the occipital, and temporal horn to the temporal. Only a thin strip along the inferolateral aspect of the frontal horn and superomedial aspect of the temporal horn segmented to the insula, constituting approximately 5% of VZ surface area, significantly lower than the other lobes (p<0.001; **Supplementary Fig. 3c).** Euclidean distance from the VZ to the insula was also significantly higher than the other lobes (p < 0.0001; **Supplementary Fig. 3d).** We then quantified the degree of obstruction (radial overlap) from subcortical structures to each lobe by sampling N=50 points from each VZ lobe segmentation at each GA **(Supplementary Fig. 3b).**

To determine the degree of obstruction, we fixed a sphere at each putative origin point in the VZ and radially projected the locations of subcortical and cortical structures on this sphere. If a subcortical structure obstructs radial paths (for radial glial cells and neural progenitors) to a cortical structure, there will be high overlap in projection space and vice versa **(Fig. 3a-f** for explanation). The lenticular nuclei blocked up to 60-70% of radial paths from the VZ to the insula. Spherical overlap of the lenticular nuclei to the Insula **(Fig. 3g)** was significantly higher than that to other lobes **(Fig. 3h** – frontal lobes). The degree of spherical overlap was significantly lower in the frontal (−26.6% [−27.3 - −25.8], p < 0.01), temporal (−30.0% [−31.2 - −29.7], p < 0.01), parietal (−30.5% [−31.2 - −29.8], p < 0.01), occipital (−30.2% [−31.0 - −29.5], p < 0.01) lobes than in the insula across gestation **(Fig. 3i).** Due to the lenticular nuclei, there were few direct radial pathways from the VZ to the insular cortex in contrast to the abundant paths to other cortices. In contrast, the caudate nucleus did not block a significant portion of paths to any lobe (at most median 10% of frontal lobe paths; **Supplementary Fig. 4a)** and the thalamus expectedly had a negligible degree of obstruction **(Supplementary Fig. 4b).**

**Fig. 3.**
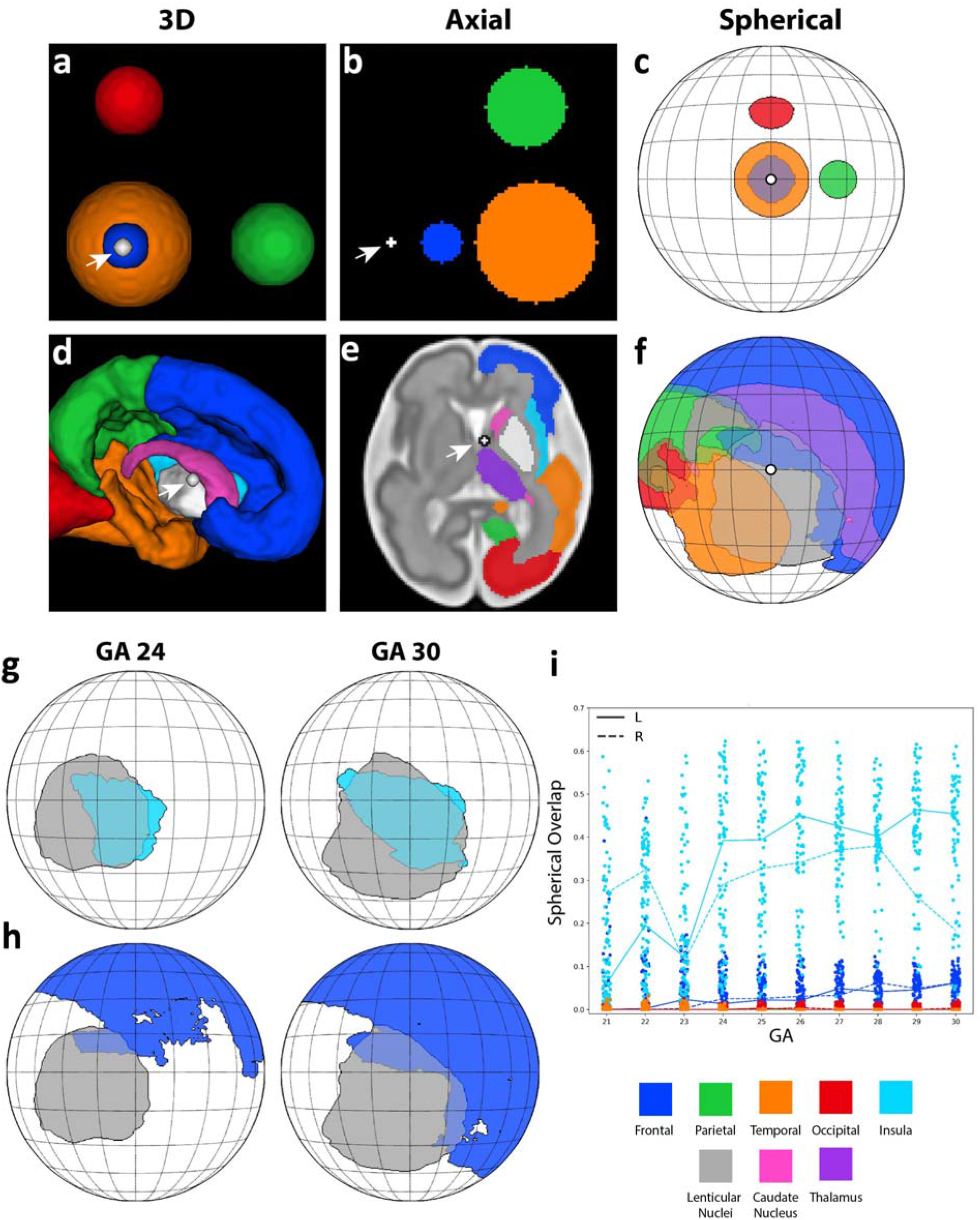
The lenticular nuclei obstruct a direct radial pathway from the ventricular zone to the insula. We developed a spherical projection analysis that determined the degree to which subcortical structures obstruct direct radial migration of radial glial cells from the ventricular zone to the cortical surface. High overlap indicates that radial pathways are obstructed. ***First row (a-c):*** Toy example using spheres. **a:** 3D visualization from the center point (*white ball and white arrow*). **b:** Axial plane – note that the red sphere is out of plane and not visualized. **c:** Spherical projection plot – the center point is at the center of the plot. Note that spherical projection warps the shape but not the solid angle taken up by a given region of interest. ***Second row (d-f):*** Example using the left side at gestational week 30. The center point is the ipsilateral foramen of Monro (*white ball and white arrow*). **d:** 3D visualization looking medial to lateral at the left hemisphere. *e:* Axial projection – note that the lenticular nuclei (gray) largely overlap the solid angle subtended by the insula **f:** Spherical projection plot. Insula and thalamus are not displayed for clarity. We determined the spherical overlap of subcortical structures with the VZ origin of each lobe. **g:** The lenticular nuclei (gray) overlap the majority of the pathways to the insula surface. *Left*: GA 24, *Right:* GA 30. **h:** The lenticular nuclei overlap a small proportion of the frontal lobe. **i:** Spherical overlap for each sample (dots) and median overlap by lobes (lines) by GA. The lenticular nuclei overlap the insula significantly more than the other lobes. Linear regression against side, GA, and lobe was significant but with moderate overall fit (R^2^ = 0.674, F(6,4842)=1672, p<0.0001). The proportion of spherical overlap was significantly lower in the frontal (−26.6% [−27.3 - −25.8], p < 0.01), temporal (−30.0% [−31.2 - −29.7], p < 0.01), parietal (−30.5% [−31.2 - −29.8], p < 0.01), occipital (−30.2% [−31.0 - −29.5], p < 0.01) lobes than in the insula. GA had a negligible effect on overlap.

The spherical projection analysis indicated that the majority of radial pathways from the VZ to the insula were obstructed by the lenticular nuclei. However, to rule out radial glia scaffolds traveling through the developing basal ganglia, we examined the DTI radiality from GA 21-30 weeks at the subplate (approximated by the gray/white matter boundary surface in the Developing Human Connectome Pipeline). DTI radiality measures the degree to which diffusion is radially oriented (e.g. perpendicular) to a given surface **(Supplementary Fig. 5)**.^30^ After 30 weeks, radiality demonstrates a steady decline due to formation of association white matter, sulcation, and other processes **(Supplementary Fig. 5d),** but prior to that, identifies the highly ordered anisotropic radial glial fascicles. Radiality at the frontal, parietal, temporal, and occipital lobes was almost 1.0 from GA 21 to 30 weeks, indicating perfect radial orientation of radial glial fascicles **(Supplementary Fig. 5b).** In contrast, radiality was significantly lower along the insula (p < 0.001), indicating an oblique direction of the glial scaffold along which progenitor cell streams could migrate **(Fig. 4e, h; Supplementary Fig. 5c).** Per the spherical projection and DTI radiality analyses, the lenticular nuclei block a direct radial path from the VZ to the insula in contrast to direct unobstructed paths to the other lobes.

**Fig. 4:**
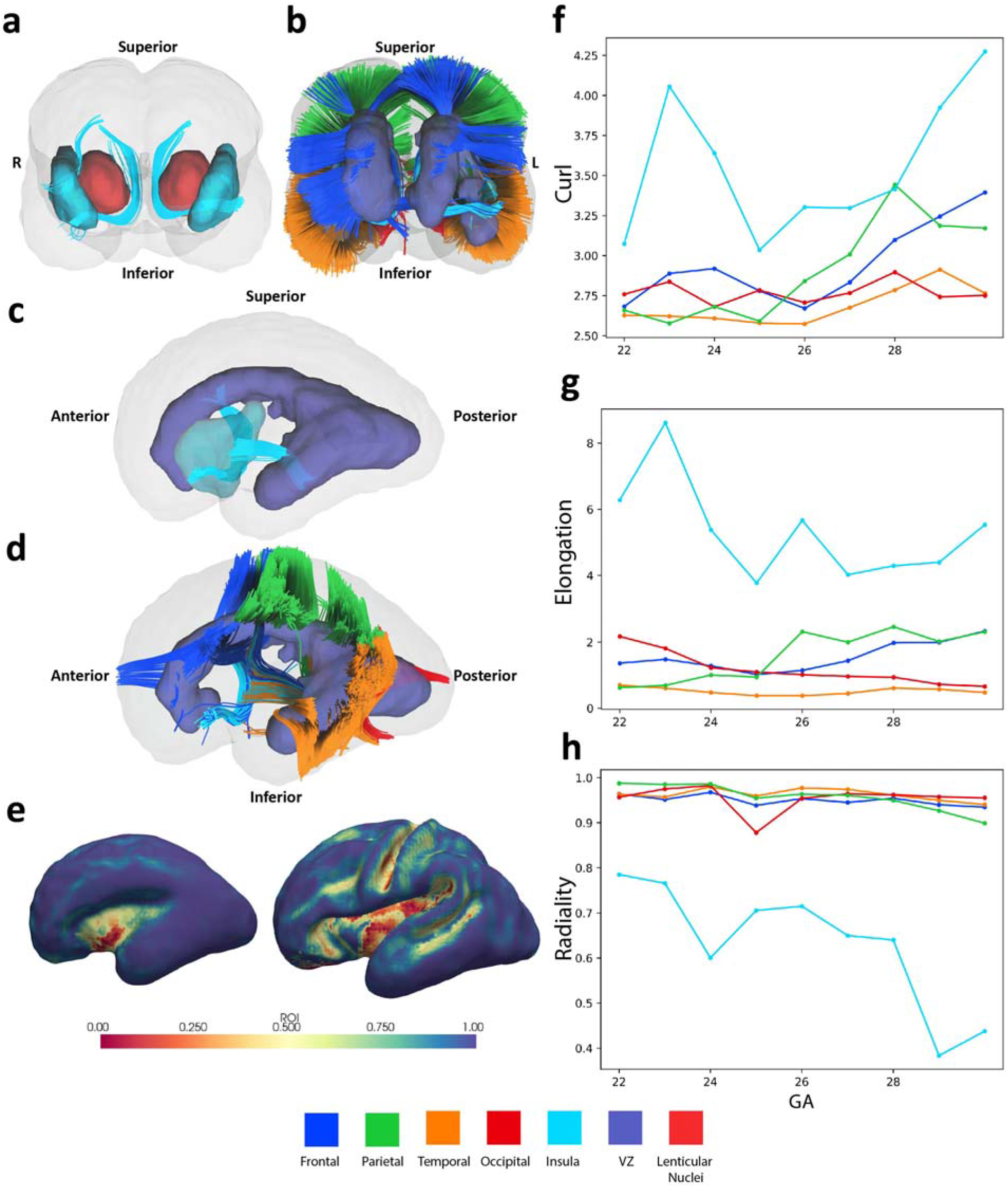
Fetal diffusion imaging demonstrates curved and oblique radial glial fascicles to insula. **a:** Diffusion tractography (anterior coronal view) demonstrating putative radial glial fascicles (light blue tractography) originating from the ventricular zone to the insula (light blue). The fascicles curve around the lenticular nuclei (red) to reach the insula. All diffusion tractography images in this Fig. are at GA 23 weeks. **b:** Diffusion tractography (anterior coronal view) with radial glial fascicles from the VZ to the frontal, temporal, parietal, and occipital lobes. Note the direct, unobstructed, orthogonal trajectory. **c:** Diffusion tractography (lateral view) of the radial glial fascicles from the VZ to insula. In addition to the fascicle curving around the lenticular nuclei, there is a second fascicle traveling obliquely from the temporal horn to the posteriorinferior insula. **d:** Diffusion tractography (lateral view) of the radial glial fascicles from the VZ to the other lobes. Again, note the direct, orthogonal pathway. **e:** Radiality^30^ at the subplate. Note the significantly lower radiality in the insula compared to the other lobes, suggesting an oblique direction for radial glial fascicles reaching the insula. *Left:* GA Week 26, *Right:* GA week 30. **f:** Diffusion tractography shape analysis^31^ - curl (total length divided by Euclidean distance between termini) by GA. Curl is significantly higher for fascicles reaching the insula than the other lobes. Linear regression against side, GA, and lobe was significant with moderate overall fit (R^2^ = 0.540, F(6,83)=18.41, p<0.0001). Mean curl was significantly smaller for fascicles going to the frontal, temporal, parietal, and occipital lobes than to the insula (p<0.001). **g:** Diffusion tractography shape analysis - elongation (total length divided by tract diameter) by GA. Elongation is significantly higher for fascicles reaching the insula than the other lobes. Linear regression against side, GA, and lobe was significant with good overall fit (R^2^ = 0.747, F(6,83)=44.71, p<0.0001). Mean elongation was significantly smaller for fascicles going to the frontal, temporal, parietal, and occipital lobes than to the insula (p<0.001). **h:** Radiality at the subplate by GA. Linear regression against side, GA, and lobe was significant with high overall fit (R^2^ = 0.838, F(6,83)=77.62, p<0.0001). Radiality was significantly higher for the frontal, temporal, parietal, and occipital lobes than the insula (p<0.001).

### The curved radial glial fascicle to the insula

The preceding analysis indicates that there is not a direct radial path to the insula. Accordingly, we used diffusion tractography to identify radial glial fascicles to the insula and compare them to those to the other lobes. Diffusion tractography identified radially oriented glial scaffolds originating in the VZ and ending in the frontal, parietal, temporal, and occipital cortices **(Fig. 4b, d).** While diffusion imaging does identify that type of fiber inducing restricted water diffusion, the radial orientation and origin of these fibers identifies them as radial glial fascicles. In contrast to the other lobes, the fibers to the insula were either curved around the lenticular nuclei **(Fig. 4a)** or traveling obliquely from the temporal VZ to the posterior insula. We quantified the degree of difference between insular radial glial fascicles and the other lobes using shape analysis.^31^ Fibers to the insula had significantly higher curl (p < 0.001; **Fig. 4f)** and elongation (p < 0.0001; **Fig. 4g),** as well as smaller diameter, shorter length, and smaller span (p < 0.001; **Supplementary Fig. 6b-d).** While mean tract fractional anisotropy was not significantly different between the insula and other lobes **(Supplementary Fig. 6a),** in isolation it is difficult to interpret the tissue composition of these tracts.

Per our analysis, a different migratory stream curving around the lenticular nucleus or traveling obliquely from the temporal VZ forms the insula in contrast to the direct radial glial fascicles to the other lobes.

## DISCUSSION

During the second and third trimesters, the cerebral cortex expands from a smooth telencephalic vesicle to a folded surface with a well-defined pattern of lobes and conserved sulci and gyri.^32,33^ While existing physical models of brain folding^14–16,34,35^ can explain the formation of an individual gyrus and sulcus, the formation of the global geometry of the brain and specific lobes remains unexplained. Moreover, recent evidence suggests that the Sylvian fissure forms by an alternative process than other sulci^20,23^, motivating the need for a better understanding of the development of the cerebrum as a whole. At the center of the C-shaped cerebrum is the insula. Buried in the depths of the Sylvian fissure, the insula has unique morphology^18,19^, cytoarchitecture^17,36^, and functional organization^37^ as compared to the other lobes. Indeed, these differences are so pronounced that almost all accounts of gyral/sulcal development specifically exclude the insula from their analyses.^15,30,35^ Radial migration models^16^ do not explain the geometry of insular gyri nor the overall configuration of the cerebrum. The anomalous nature of insular development suggest that fundamentally different processes are involved.

This study quantitively establishes that the insula has different morphology than the other lobes in adults **(Fig. 1)** and demonstrate that these differences emerge during fetal development **(Fig. 2).** Specifically, the insular cortex expands at a slower rate than any other brain lobe and has a unique gyral arrangement and morphology, different than the resulted geometry from radial migration models^12,14–16,34,35^. Jacobian analysis establishes that the differences in volumetric growth were most pronounced between the insula (low growth) and the surrounding operculae (high growth) **(Fig. 2g).** This stark difference in volumetric growth closes the Sylvian fissure and brings the anterior temporal into proximity with the basal frontal lobe, establishing the C-shape of the cerebrum.

We then identify a novel mechanism by which these differences emerge. Using a novel spherical projection method **(Fig. 3),** we describe how the lenticular nuclei directly obstruct a radial pathway between the ventricular zone and most of the insular cortex throughout gestation, an obstacle that is not observed for the other lobes. Finally, using fetal diffusion imaging **(Fig. 4),** we demonstrate that a curved fascicle travels *around* the lenticular nuclei to the insula and another tangentially traveling fascicle travels to the posterior insula from the temporal VZ **(Fig. 5).** Fetal diffusion imaging is still in its infancy ^30^, but our regional comparative analysis demonstrates that glial scaffolding for migrating progenitor cells forms around but not through the lenticular nuclei to the insula, a new finding that aligns with other results presented here.

**Fig. 5:**
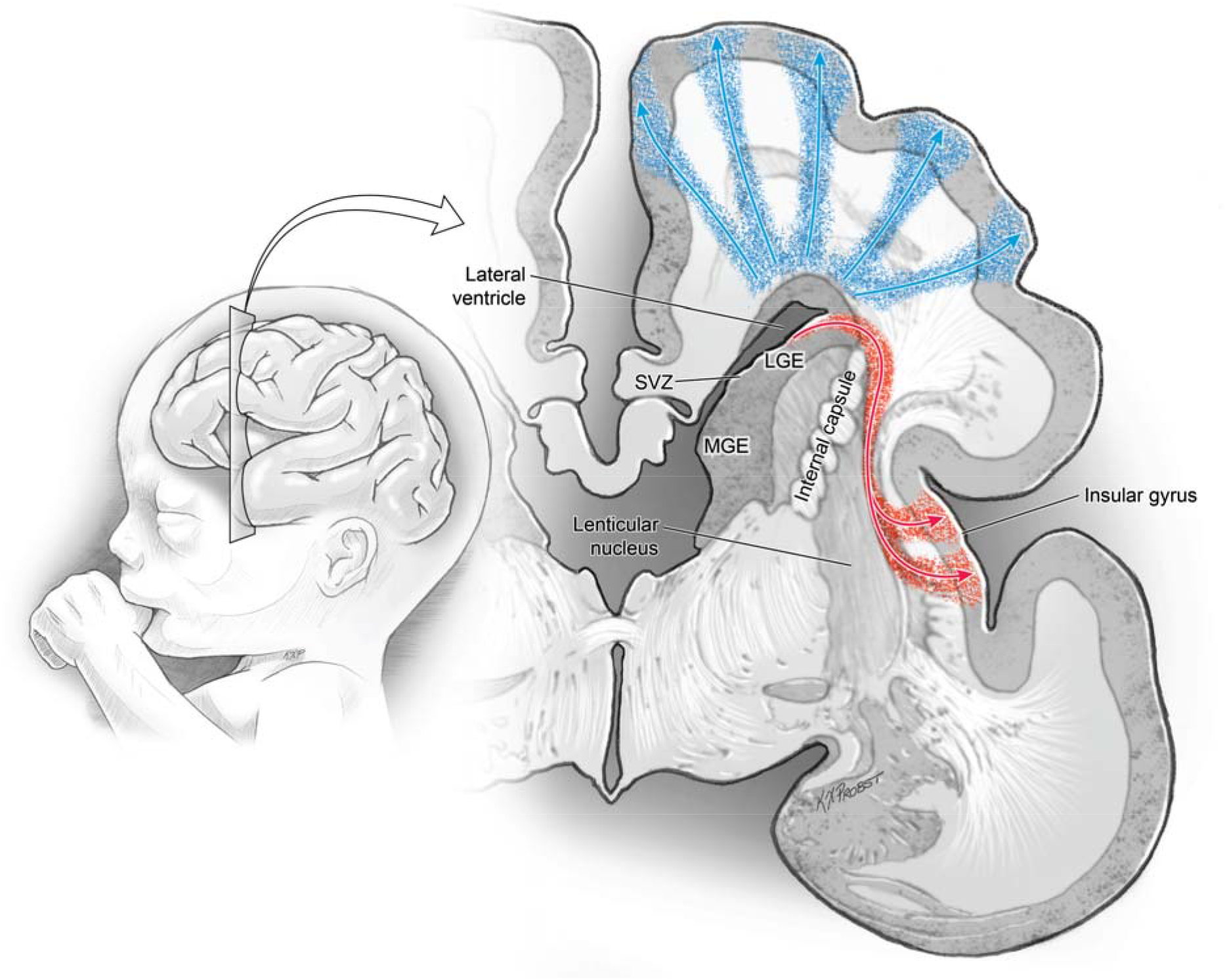
A novel mechanism for insular development and the formation of the global geometry of the cerebrum. The lobes of the cerebral convexity (frontal, parietal, temporal, occipital) form via direct radial migration along radial glial scaffolds (blue). In contrast, a direct radial pathway from the ventricular zone to the insula is obstructed by the lenticular nuclei. Consequently, neural progenitor cells travel along a curved and oblique scaffold around the lenticular nuclei to the insula. Accordingly, insular gyri have an altered geometry, and the insula expands at a significantly lower rate than the other lobes. This relative overgrowth of the operculae closes the Sylvian fissure, buries the insula, and defines the global configuration of the cerebral lobes.

Both evolutionary and cytoarchitectonic evidence support a unique development of the insula. For instance, despite having an overall gyrencephalic cortex on the cerebral convexity, macaques have lissencephalic insula, indicating that different mechanisms gyrify both cortices.^38^ Cytoarchitectonically, the insula itself is arranged into horizontal/oblique stripes of agranular, dysgranular, and granular cortex that run transverse to the direction of the insular gyri. In contrast to several locations on the convexity, these cytoarchitectonic regions do not respect sulcal boundaries and instead seem to emanate from the limen insula.^39,40^ The insula subserves primary interoceptive functions (in addition to others), and the transfer of information from granular to dysgranular to agranular cortices may support this function.^17^

Finally, the spectrum of malformations of cortical development again supports a unique development process in the insula. The stereotypic finding of the insula buried in the Sylvian fissure of a C-shaped cerebrum is conserved in a wide variety of malformations including lissencephaly and polymicrogyria, in which all other cortical folds are disrupted.^41^ In contrast, malformations of the telencephalic flexure – e.g. the bending of the cerebrum into the C-shape and formation of the Sylvian fissure, such as the holoprosencephaly spectrum, do not display an insula.^42^ This highlights the key role of the insula in organizing the telencephalon. Consequently, our findings explain not only the architectural and functional differences in the insula but also the role of this mechanism in forming the global geometry of the cerebrum.

Our results indicate that the brain does not grow homogeneously. The central core and insular cortex are developmental outliers. The opercula receive radial migration from the ventricular zone and grow at a faster rate than the neighboring insula and central core, resulting in deep, convoluted gyri. In contrast, a curved and tangential glial fascicle travels from the VZ, around the lenticular zone, to the insula **(Fig. 5).** Accordingly, the insular folds have little in common with their equivalents on the cerebral convexity and the insula and the central core expand a much slower rate. The rapidly expanding operculae overgrow the insula and close the Sylvian fissure, analogous to the way a muffin top overgrows the stump. As we have shown before, the formation of the Sylvian fissure follows different principles compared to the development of any other brain fold and that singularity explains why arteries do not cross the Sylvian fissure.^20,23,43^ Those findings are confirmed by the results of this paper.

The putative origin of insular progenitors indicated by this analysis **(Fig. 5; Supplementary Fig. 3A)** corresponds to the pallial/subpallial boundary, the interface zone between the ganglionic eminence, which originates tangentially migrating neurons, and the ventricular zone, which originates radially migrating cortical progenitors.^2,44^ While tangential migration from the ganglionic eminences themselves has been identified ^6,45,46^, Gonzalez-Arnay et al. hypothesized that this area of the ventricular zone between the pallium and the subpallium generates insular progenitors that also migrate tangentially.^47^ Our findings confirm this hypothesis. The pallial-subpallial boundary is well positioned for progenitor migration around the basal ganglia. Furthermore, cells at this boundary are genetically distinct from the pallium which may additionally contribute to morphological differences in gyri.^48^ Finally, the subpallium is a source of numerous structures with strong reciprocal connections with the insula^49^ and may exhibit a strong influence on cortical development including the insula.^48^ The close proximity of the pallial-subpallial boundary to putative insular progenitors may facilitate this process.

Interestingly, our findings were observed by previous authors but not fully explored. For instance, Hill and colleagues observed significantly slower surface area expansion in the insula during post-natal development.^35^ Similarly, Garcia et al. observed relatively slower growth in the insula in pre-term infants but did not provide a mechanistic explanation.^34,50^ Of the several present theories of cortical gyrification^51^, none specifically address the morphological differences in the insula nor the role its low growth plays in forming the global arrangement of the cerebrum. In fact, several of the keystone papers in the theories of axonal tension^15^ and constrained tangential cortical expansion^14,16^ either do not address the insula at all or predict anatomically incorrect patterns of gyration in the insula. Similarly, past attempts to identify universal scaling laws relating cortical thickness and cortical surface area specifically exclude the insula.^52,53^ Given our findings of a putative pallial-subpallial origin of insular progenitors, cellular specification models likely can shed light on insular development^29,32^, but none so far have specifically addressed this unique population of insular progenitors. Consequently, this report explains the unique geometric properties of the insula with a novel model of insular cortical development.

As longitudinal collection of fetal brain specimens from the same individual is logically impossible, fetal MR imaging is an optimal tool to study the still open problem of human brain folding. However, further work in fetal brain specimens may further validate these results.

## CONCLUSION

We identify a new model of insular development in contrast to direct radial migration. Unlike the other lobes, the insula forms from curved and tangential migratory streams that travel around the lenticular nuclei and from the temporal ventricular zone. This novel mechanism results in both the altered morphology and the relative slower growth of the insula. Accordingly, the growth of the surrounding frontal, temporal, and parietal operculae, driven by direct radial migration, outstrips that of the insula and creates the characteristic C-shaped form of the cerebrum around the Sylvian fissure.

## METHODS

### Data Sources

#### Fetal

Fetal anatomic data was obtained from the spatiotemporal atlas published by Gholipour et al.^28^ This atlas was reconstructed from weekly fetal T2-weighted MRI images of 81 healthy fetuses at a GA between 19 and 39 weeks. A full description of the Atlas and its content can be accessed at http://crl.med.harvard.edu/research/fetal_brain_atlas/. The authors obtained images using repeated T2-weighted half-Fourier acquisition single shot fast spine echo series (TR=1400-2000ms, TE=100-120ms, variable FOV, 2mm slice thickness, 0.9-1.1mm in plane resolution). Images were preprocessed with motion and bias field correction and superresolution volumetric reconstruction. The atlas was then constructed from volumetric reconstructions utilizing kernel regression over time and symmetric diffeomorphic deformable registration in space.^54^ The final atlas resolution was isotropic 1mm^3^ obtained using robust super-resolution volume reconstruction. The atlas segmentation was initialized using a previous segmentation of cortical and subcortical regions developed by Gousias et al.^55,56^, manually edited, and refined extensively for all gestational ages in the process described in ^28^.

Fetal diffusion data was obtained from the atlas published by Khan et al.^57^ and available at http://crl.med.harvard.edu/research/fetal_brain_atlas/. Diffusion tensors were not available in the online repository and obtained directly from the authors of that study. Full details are available from the original publication. Briefly, the authors obtained 67 fetal MRI from 60 subjects ranging from 21 weeks to 38 weeks on a 3T Siemens Skyra MRI. with 16-channel body matrix and spine coils. Each DTI acquisition consisted of 1-2 b=0 s/mm^2^ images and 12 b=500 s/mm^2^ over 2-8 scans in one of three orthogonal planes to the fetal head. Acquisition parameters were TR=3000-4000ms, TE=60ms, 2mm x 2mm resolution with 2-4 mm slice thickness. Structural images were reconstructed per the authors’ pipeline. Following this, B=0 and B=500 images were registered to anatomic space using a robust Kalman-filter based slice-to-volume registration that accounted for subject motion during acquisition^58^. The authors then calculated diffusion tensors in anatomic space. These tensors were then used to calculate an unbiased spatiotemporal diffusion tensor atlas integrating kernel-regression in age with a diffeomorphic tensor-to-tensor registration. The diffusion tensors from the atlas at each GA from 22-30 weeks were used for this analysis.

#### Adult

We randomly selected 107 adult patients (54.2% female) of median age 28 (range 22-35) from the structural data in the Human Connectome Project HCP 900 subjects data release.^24^ Specific subject IDs are listed in **Supplementary Table** 1. Further details and data can be seen at https://www.humanconnectome.org/study/hcp-young-adult/document/900-subiects-data-release. The details for acquisition are reported on that website. Briefly, T1 and T2 weighted imaging was obtained on Siemens 3T platforms using a 32-channel head coil. Each subject had T1 and T2-weighted imaging as well as CIFTI cortical/surface segmentation in native and atlas space. For this report, all analysis was performed in native space. The cortical segmentation with the Desikan-Killiany atlas^59^ was used for this analysis. Specific gyri and sulci segmented in the Desikan-Killiany atlas were mapped to frontal, temporal, parietal, and occipital lobes as well as the insula per the instructions at https://surfer.nmr.mgh.harvard.edu/fswiki/CorticalParcellation.

### Volumetric segmentation and analysis

We performed volumetric segmentation in the fetal cohort by lobe (frontal lobe, temporal lobe, parietal lobe, and occipital lobe as well as the insula). Each non-insular lobe is associated with superficial and deep white matter which constitutes most of the volume of said lobe. In contrast, the insula does not have a deep white matter component and only a thin layer of subcortical white matter (the extreme capsule). Consequently, per the analysis of Türe et al. and Ribas et al.^18,19^ we identified the insula as the cortex associated with the central core of the brain. The central core is an anatomic concept consisting of the deep telencephalic structures of the cerebrum, specifically the caudate nucleus, lenticular nuclei (putamen, globus pallidus externus and internus), and the internal capsule. Ribas describes the thalamus as part of the central core. We calculated the volume of cortical and subcortical structures by multiplying the total number of voxels segmented to each structure multiplied by the volume of each voxel (1mm^3^). Unless otherwise specified, results are presented for the left hemisphere as we uniformly did not find a significant difference between the right and left hemispheres. Volumetric segmentation and analysis were performed using the *NumPy, ANTsPy*, and *NiBabel* packages in Python (Python 3.7)

### Surface segmentation and analysis

The adult data was provided with CIFTI maps of the cortical surface at the pial surface, midthickness, and gray-white matter interface as well as maps of cortical thickness, sulcal depth, mean curvature. Adult segmentations were mapped to lobes as described above. Fetal brain surface segmentation was performed using the Developing Human Connectome Pipeline (dHCP) reported by Makropoulos et al.^60^ on our fetal anatomic dataset. The full pipeline is described at http://github.com/BioMedIA/dhcp-structural-pipeline. The atlas for the structural pipeline was the same one as above (Gousias 2012)^56^ and was remapped to frontal/temporal/parietal/occipital/insula manually. dHCP results were manually inspected for artefacts and anatomic validity. Similar to the adult dataset, the dHCP pipeline generated maps of cortical thickness, sulcal depth, and mean curvature. In addition for both adult and fetal data, we generated maps of Gaussian (intrinsic) curvature using Connectome Workbench and fractal dimension, using the dilation method described by Madan et al.^25^, coded in Python (Python 3.7).

In both adults and fetal datasets, surface area was calculated using the midthickness segmentation by lobe. We calculated mean cortical thickness, mean curvature, Gaussian curvature, cortical thickness, and fractal dimension by lobe for each adult subject and fetal GA. For sulcal depth, the HCP/dHCP algorithms calculate a higher sulcal depth for all areas of the insula as it is buried in the depths of the Sylvian fissure. However, our primary interest was the *within* lobe sulcal depth – i.e. the distance from the deepest sulcus to the highest gyral crown in each lobe. Consequently, we defined a new metric, sulcal depth range (SDR), defined as the difference between the lowest and highest values of sulcal depth per lobe.

Fetal data was plotted as a line plot by time while adult data (due to the lack of a temporal variable) was plotted as box and whisker plots using a combination of tools from the Python packages *matplotlib, seaborn*, and *plotnine*. Unless otherwise specified, results are presented for the left hemisphere only as we uniformly did not find a significant difference between the right and left hemispheres.

### Adult and fetal data analysis and visualization

#### Linear and non-linear regression

For each morphometric variable (volume – fetal only, surface area, cortical thickness, sulcal depth range, mean curvature, Gaussian curvature, fractal dimension), we performed multiple regression to identify a lobe-specific effect. For the fetal data, we regressed each metric against lobe, controlling for gestational age and laterality. Similarly, for adult data, we regressed each metric against lobe, controlling for laterality. Significance of the multiple regression was defined as a prespecified level of p < 0.01. In the adult dataset only, metrics that had significant effect of lobe, *post hoc* Mann-Whitney testing was performed between each non-insular lobe and the insula, with Bonferroni correction to avoid issues with multiple comparisons. Statistics were performed using the *pandas* and stats models package of Python as well as R 4.2.0 (R Foundation for Statistical Computing) (with *rpy2* bridge to Python).

In addition, to quantify the trajectory of fetal brain development by GA, we performed nonlinear regression of each metric using a logistic, exponential, and a linear model **(Supplementary Table 2).** Best model fit was determined using Akaike information criteria (AIC). The *optimize* package in *SciPy* was utilized for curve fitting.

#### Data visualization

Given the high dimension of our acquired morphometric database, we performed UMAP (Uniform Manifold Approximation and Projection) on the fetal and adult datasets.^61^ UMAP allows for projection of data into two-dimensions to visualize clustering and may better preserve relational information than comparable algorithms. For the adult dataset, UMAP parameters were: number of neighbors = 20, minimum distance = 0.7, seed = 281. For fetal data, parameters were: number of neighbors = 75, minimum distance = 0.4, seed = 74.

### Ventricular zone spherical overlap analysis

#### Ventricular zone segmentation by lobe

The CRL fetal brain MRI atlas contains a pre-specified segmentation of the ventricular zone (VZ) by through gestational age 30 weeks. We hypothesized that each point on the cortical surface originated from the closest point in the VZ. Consequently, we segmented the VZ to each lobe as follows. For each point in the VZ we identified the closest point on the pial surface using a K-D tree-based approach. The segmentation of each pial point (e.g frontal, insular, etc.) was assigned to the VZ point. We computed the percentage of VZ area by lobe and the mean distance from each lobe to the VZ. Statistical analysis for this data was performed similarly to morphometric data (as described above). We then randomly sampled 50 points from each VZ segmentation and each GA for spherical projection analysis. This analysis was performed using the *SciPy, NumPy*, and *PyVista* packages in Python.

#### Spherical Overlap

For this analysis we developed a novel spherical projection approach to determine the degree to which subcortical structures (e.g. lenticular nuclei, caudate nucleus, and thalamus) obstruct a direct radial path from the VZ to the cortical surface. To achieve this, we re-expressed the cortical/subcortical segmentation (described above) in spherical coordinates with each sampled VZ point as the origin. Subsequently, we projected this data along each radial vector, resulting in a projection of segmentation data on a sphere centered at each sampled VZ point. If subcortical structure highly overlapped with a cortical structure, this meant that direct radial paths from the VZ to the cortex were obstructed by said structure. See **Fig. 3a-f** for further details. We computed spherical overlap at each of the 50 sampled points for each lobe against 3 subcortical structures (lenticular nuclei, caudate nucleus, and thalamus) at each gestational age. Spherical overlap was calculated as the steradians of overlap between subcortical structure and cortex of each lobe, divided by the steradians of the given lobe (unitless). Spherical overlap for each subcortical structure was regressed against lobe, controlling for laterality and GA, using the statistical parameters described above. Spherical projection maps were plotted using the *basemap* package in Python.

### Diffusion analysis

#### Radiality

Radiality at the gray-white matter junction was computed using the approach described by Eaton-Rosen et al.^30^ Briefly, we computed the principal direction of diffusion (using DTI-TK) and the surface normal (using *PyVista)* at each point in the gray-white matter junction surface. Eaton-Rosen et al. defined the *radiality* as the normalized dot product of these two vectors. See **Supplementary Fig. 5** for further details. We computed the radiality for each point in each lobe at each GA from 22-33.

#### Tractography

Diffusion tensors were obtained from the authors of Khan et al. 2019.^57^ These tensors (by GA) were imported into DSI Studio.^62^ The fetal atlas was warped into DTI space utilizing affine registration with FLIRT.^63^ Inclusion/exclusion regions of interest (ROI) were obtained from the fetal atlas. A midline region of avoidance was hand-drawn for each GA. The deterministic fiber streamline tracking algorithm described by Yeh et al.^62^ was used. The cortical segmentation of each lobe was an ROI and the *unsegmented* (whole) VZ was specified to be an end zone (as to not bias the results with our computed VZ segmentation). The midline region of avoidance was used to avoid spurious tracts crossing midline. The fractional anisotropy threshold was 0.05. The angular threshold was 45 degrees. The step size was randomly selected from 0.5 voxel to 1.5 voxels. Tracks with length shorter than 5 or longer than 400 mm were discarded. A total of one million seeds were placed. Once tractography was completed, tracts were inspected for artefacts and for anatomic validity.

In order to perform quantitative analysis of each developmental tract bundle, we used the shape analysis tools in DSI Studio as described by Yeh et al.^31^ Specifically, we calculated tract curl, elongation, span, length, diameter, and mean fractional anisotropy. See the reference for specific descriptions of each parameter. We compared each parameter by lobe using multiple linear regression, controlling for GA and laterality using the statistical approach described above.

## Data Availability

This analysis was primarily performed in Python (Python 3.7) with a small contribution of R (R 4.2.0) using the packages described above. Additional visualization was performed using ITK-SNAP^64^ as well as Connectome Viewer. Other analysis was performed in specific, publicly available toolboxes as described in each individual section. Python and R code is publicly available for review at http://github.com/mallelaan/brain_development.

Fetal data is publicly available for download from http://crl.med.harvard.edu/research/fetal_brain_atlas/. Diffusion tensors can be obtained from the authors of Khan et al. upon request. Adult data can be obtained from https://www.humanconnectome.org/study/hcp-young-adult/document/900-subjects-data-release using the subject IDs specified in **Supplementary Table 1.**

## Acknowledgements

Data were provided [in part] by the Human Connectome Project, WU-Minn Consortium (Principal Investigators: David Van Essen and Kamil Ugurbil; 1U54MH091657) funded by the 16 NIH Institutes and Centers that support the NIH Blueprint for Neuroscience Research; and by the McDonnell Center for Systems Neuroscience at Washington University.”

In addition, the authors would like to thank the authors of Gholipour et al.^28^ and Khan et al.^57^ for graciously sharing their atlases online and for providing diffusion tensors.

## Funding

American Association of Neurological Surgeons – Van Wagenen Fellowship and Grant (EG) Burroughs Wellcome Fund Physician-Scientist Institutional Award (ANM)

National Institutes of Health 1F32DC020644-01 (ANM)

National Institutes of Health R01 EB013248 (AG, SKW)

National Institutes of Health R01 NS106030 (AG, SKW)

National Institutes of Health R01 EB031849 (AG, SKW)

National Institutes of Health R01 EB032366 (AG, SKW)

National Institutes of Health R01 HD109395 (AG, SKW)

National Institutes of Health S10 OD0250111 (AG, SKW) McKnight Foundation – Technological Innovations in Neuroscience (AG, SKW)

The content of this publication is solely the responsibility of the authors and does not necessarily represent the official views of the NIH, McKnight Foundation, American Association of Neurological Surgeons, or the Burroughs Wellcome Fund.

## Author Contributions

A.M. and E.G. conceived the project. A.G. and S.K.W. additionally developed analyses and provided data. A.M. and H.D. analyzed the data. A.M., H.D., and E.G. wrote the manuscript. A.M., E.G., H.D., A.G., and S.W. critically edited the manuscript. E.G. supervised the project.

## Competing Interests

The authors declare no competing interests

## Notes

https://github.com/mallelaan/brain_development/

### Competing Interest Statement

The authors have declared no competing interest.

### Summary of Updates

Added new figure, edited discussion and introduction

