## Supplementary figures and images for "Heterogeneous migration of neuronal progenitors to the insula shapes the human brain"

### Supplementary Figure 1

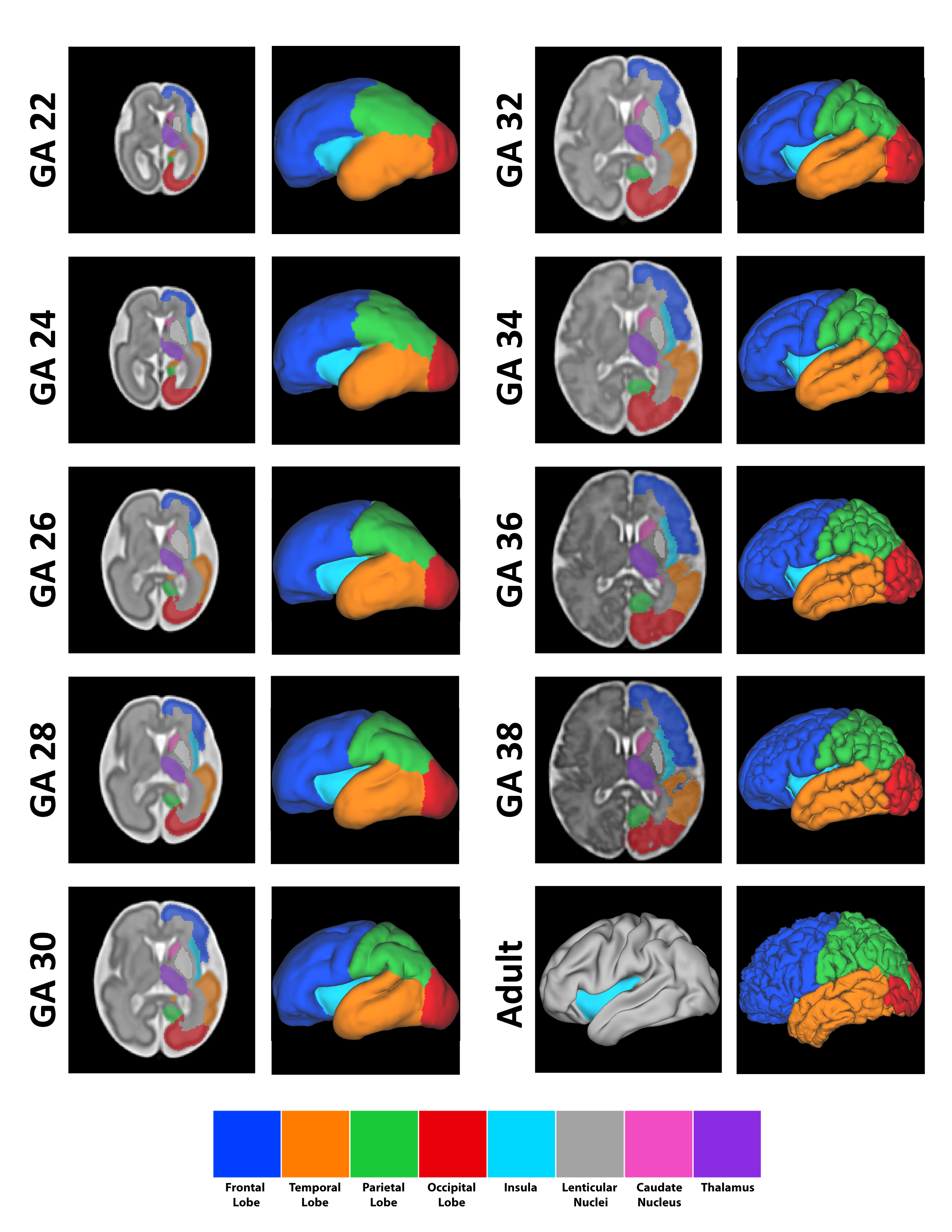

### Supplementary Figure 2

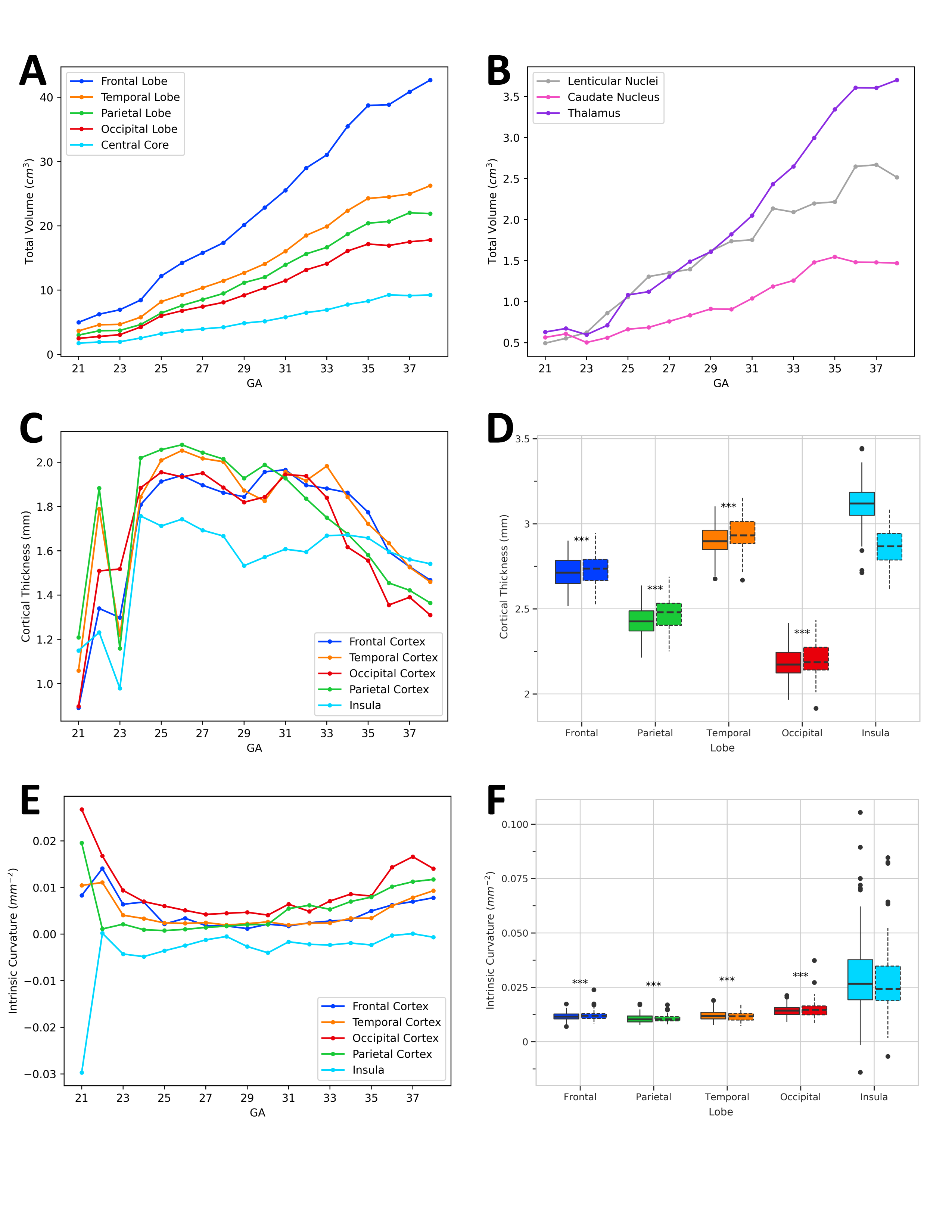

### Supplementary Figure 3

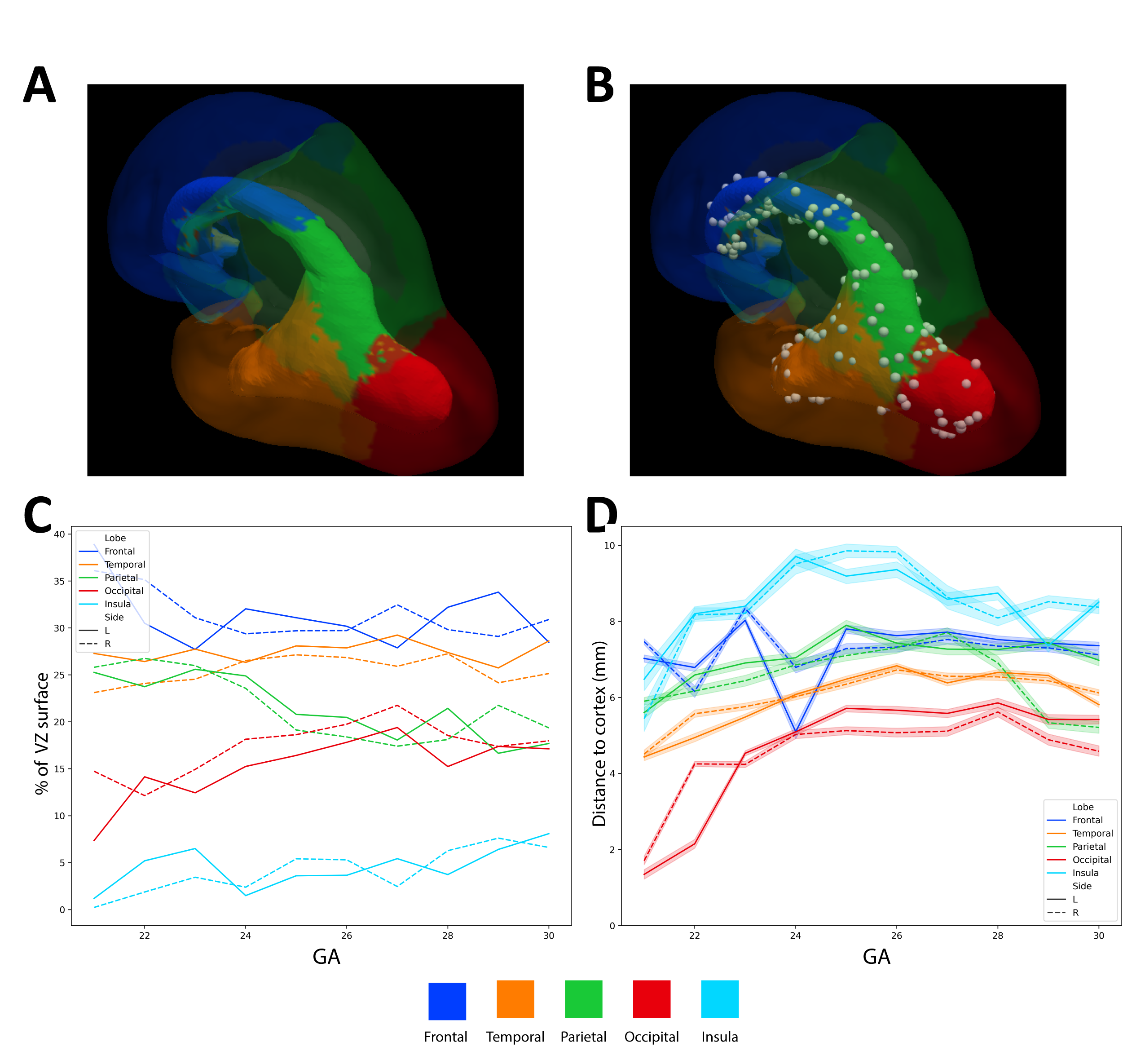

### Supplementary Figure 4

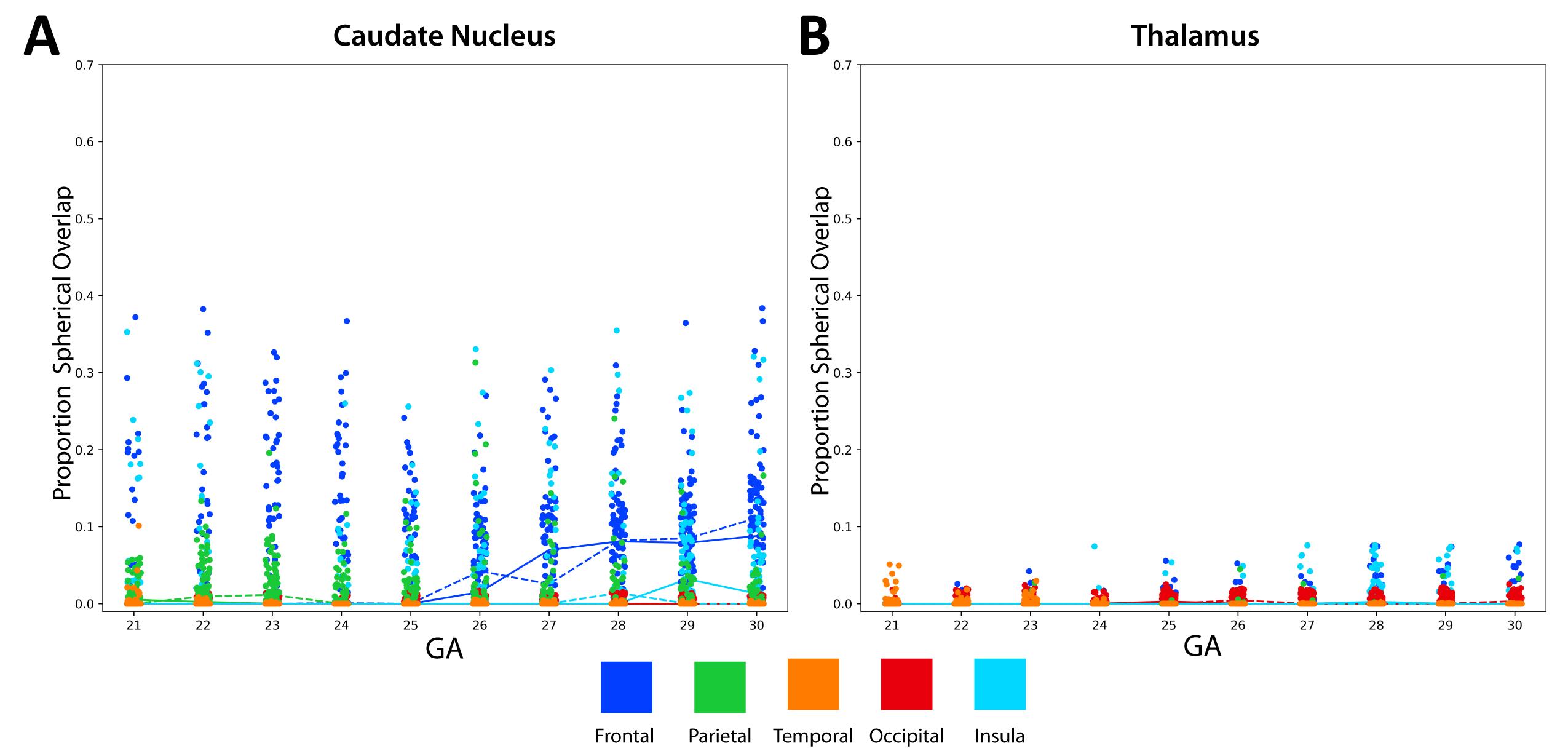

### Supplementary Figure 5

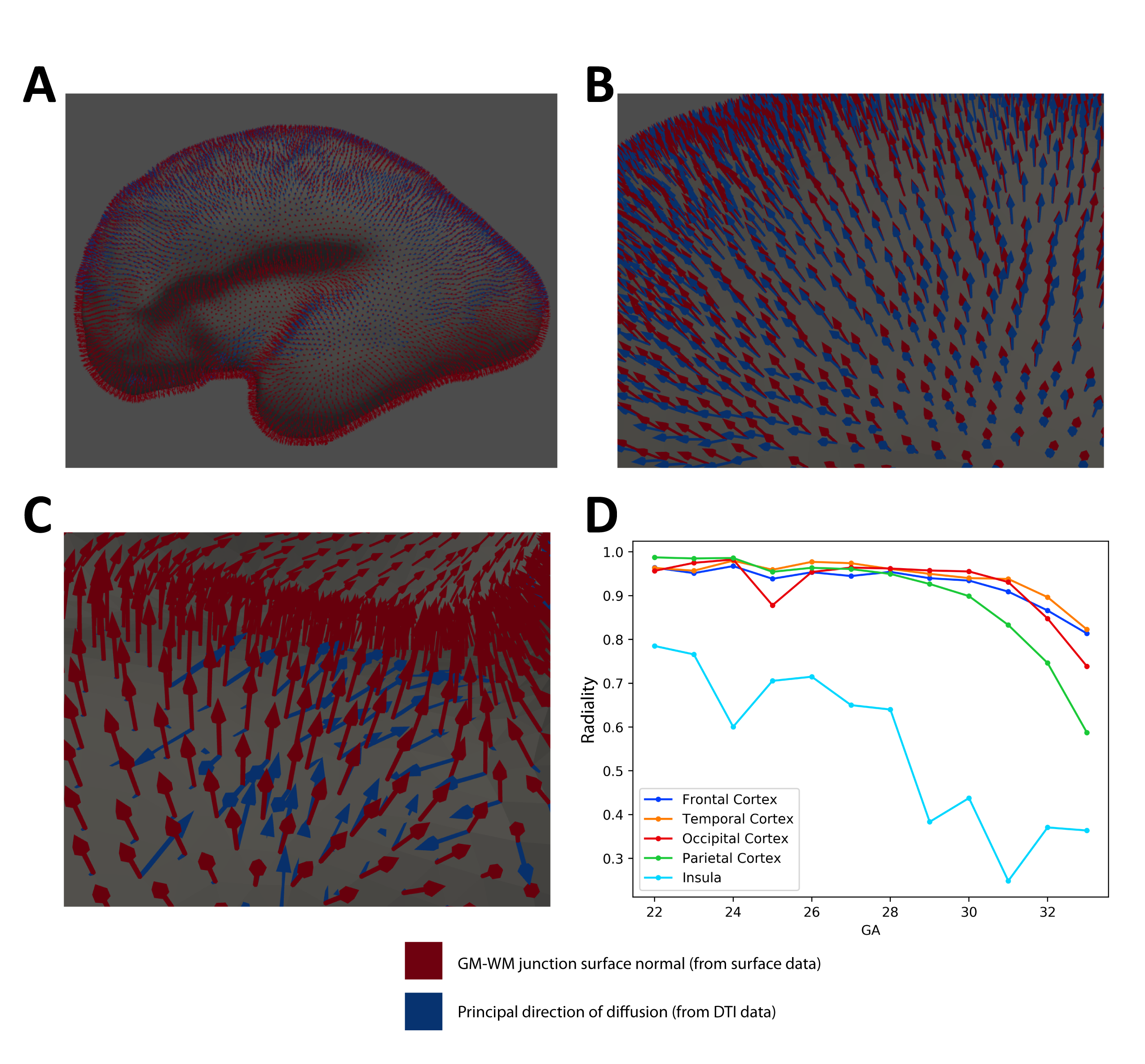

### Supplementary Figure 6

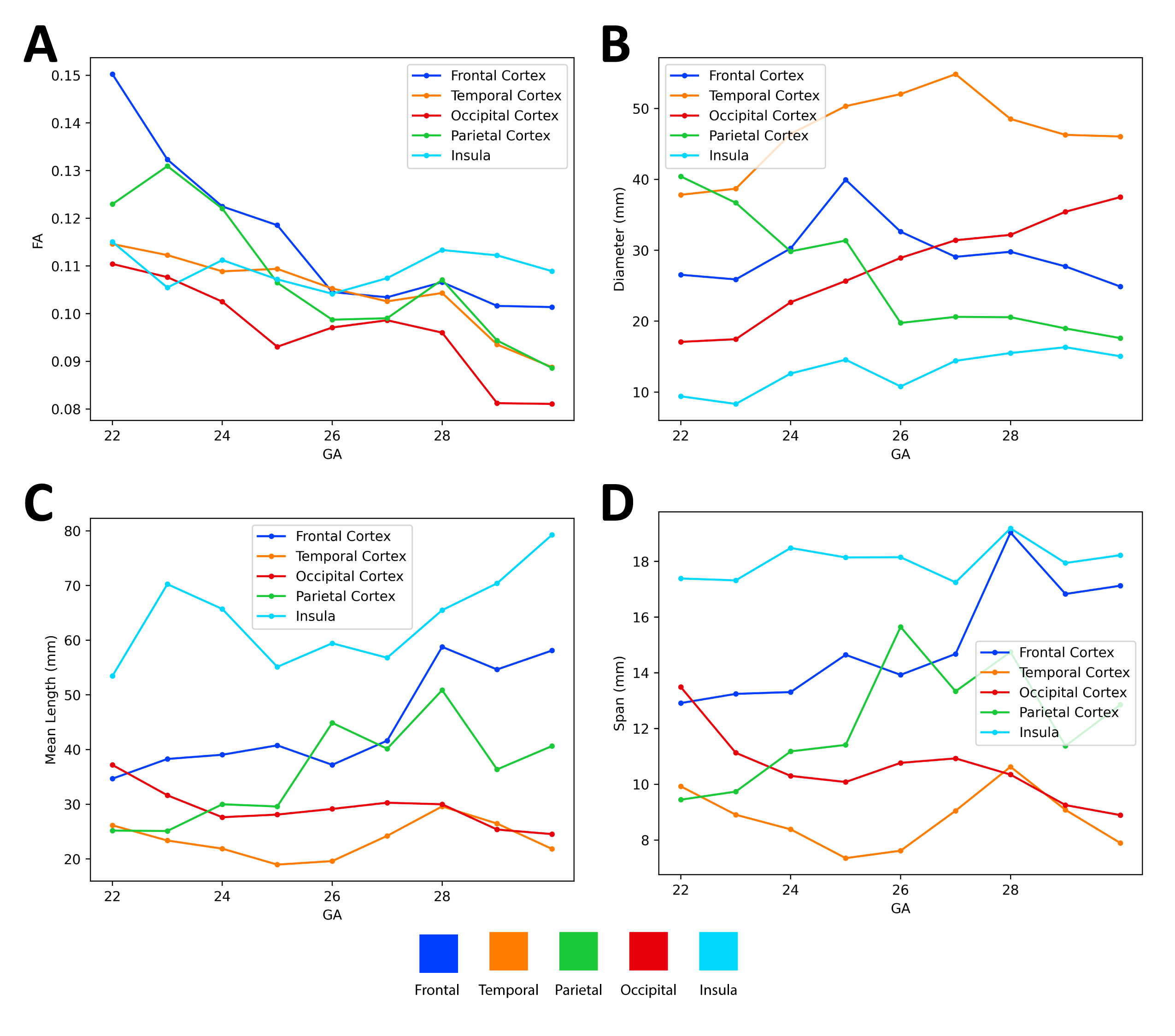
