## Supplementary Materials - Methods + Supplementary Figures/Tables for "Heterogeneous migration of neuronal progenitors to the insula shapes the human brain"

In addition, to quantify the trajectory of fetal brain development by GA, we performed non-linear regression of each metric using a logistic, exponential, and a linear model (**Table S2**). Best model fit was determined using Akaike information criteria (AIC). The *optimize* package in *SciPy* was utilized for curve fitting.

***Diffusion analysis***

*Radiality*

Radiality at the gray-white matter junction was computed using the approach described by Eaton-Rosen et al.(*30*) Briefly, we computed the principal direction of diffusion (using DTI-TK) and the surface normal (using *PyVista*) at each point in the gray-white matter junction surface. Eaton-Rosen et al. defined the *radiality* as the normalized dot product of these two vectors. See **Figure S5** for further details. We computed the radiality for each point in each lobe at each GA from 22-33. To facilitate additional comparison with Eaton-Rosen 2017 we computed linear fits (intercept and slope) for each lobe (**Table S1**),

Fetal data is publicly available for download from <http://crl.med.harvard.edu/research/fetal_brain_atlas/>. Diffusion tensors can be obtained from the authors of Khan et al. upon request. Adult data can be obtained from <https://www.humanconnectome.org/study/hcp-young-adult/document/900-subjects-data-release> using the subject IDs specified in **Table S1**.

**Supplementary Figures**

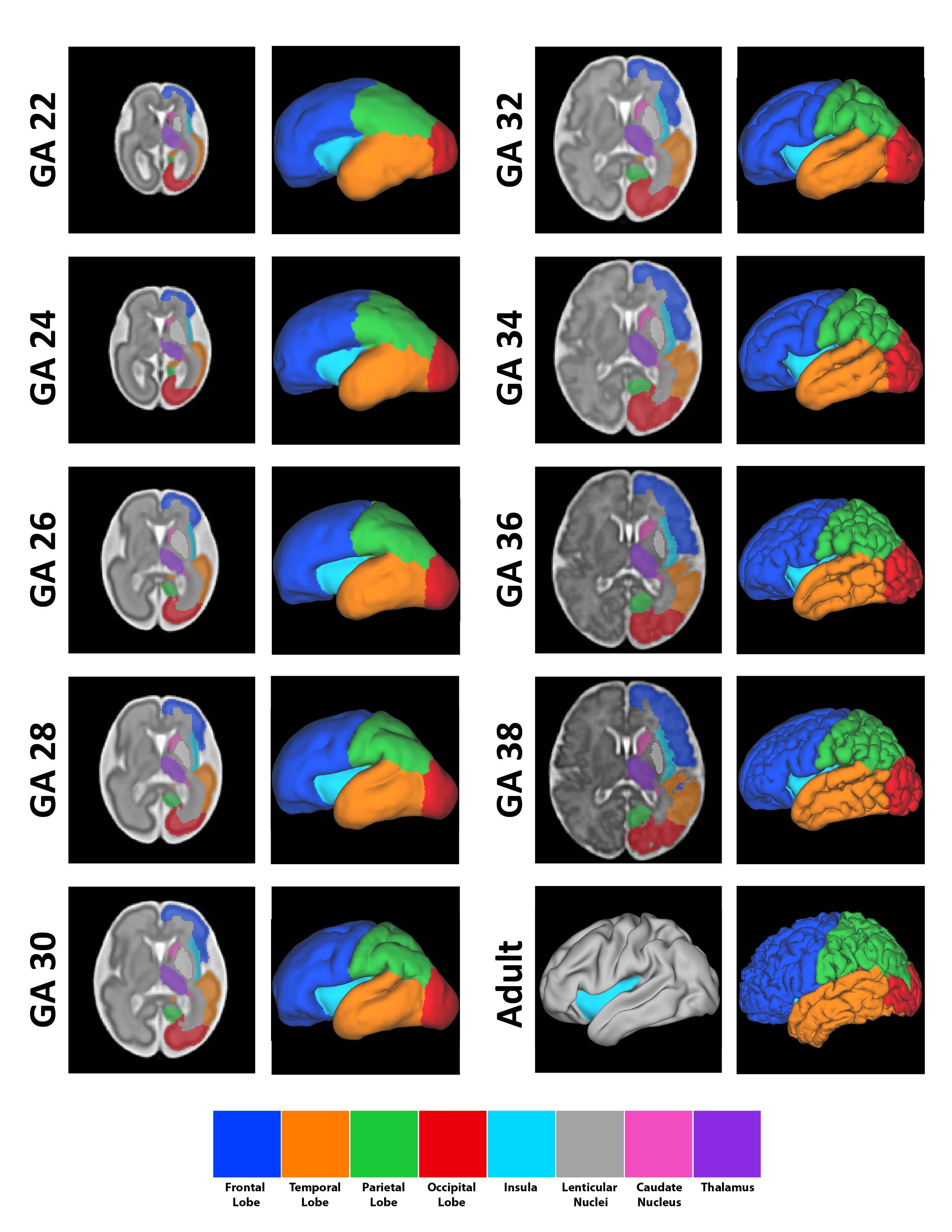

**Figure S1:** **Development of cerebral lobes, insula, and central core by gestational age (GA).** *Left side of columns* – axial image T2-weighted fetal atlas MRI (at the level of foramen of Monro) demonstrating segmentation of cortical surfaces and central core. *Right side of columns* – left lateral pial surface. Note the progressive formation of gyri/sulci in the frontal, temporal, parietal, and occipital surfaces in contrast to the smooth surface of the insula. The first sulcation of the insula (central sulcus of the insula) is noted at GA 35-36. The operculae progressively cover the insula throughout gestation until it is completely covered in adulthood. *Adult (right column, bottom row)* – *Right* *panel*: Adult pial surface. Note that the insula is covered. *Left panel*: Inflated cortical surface, demonstrating the relatively flat insula hidden under the opercula

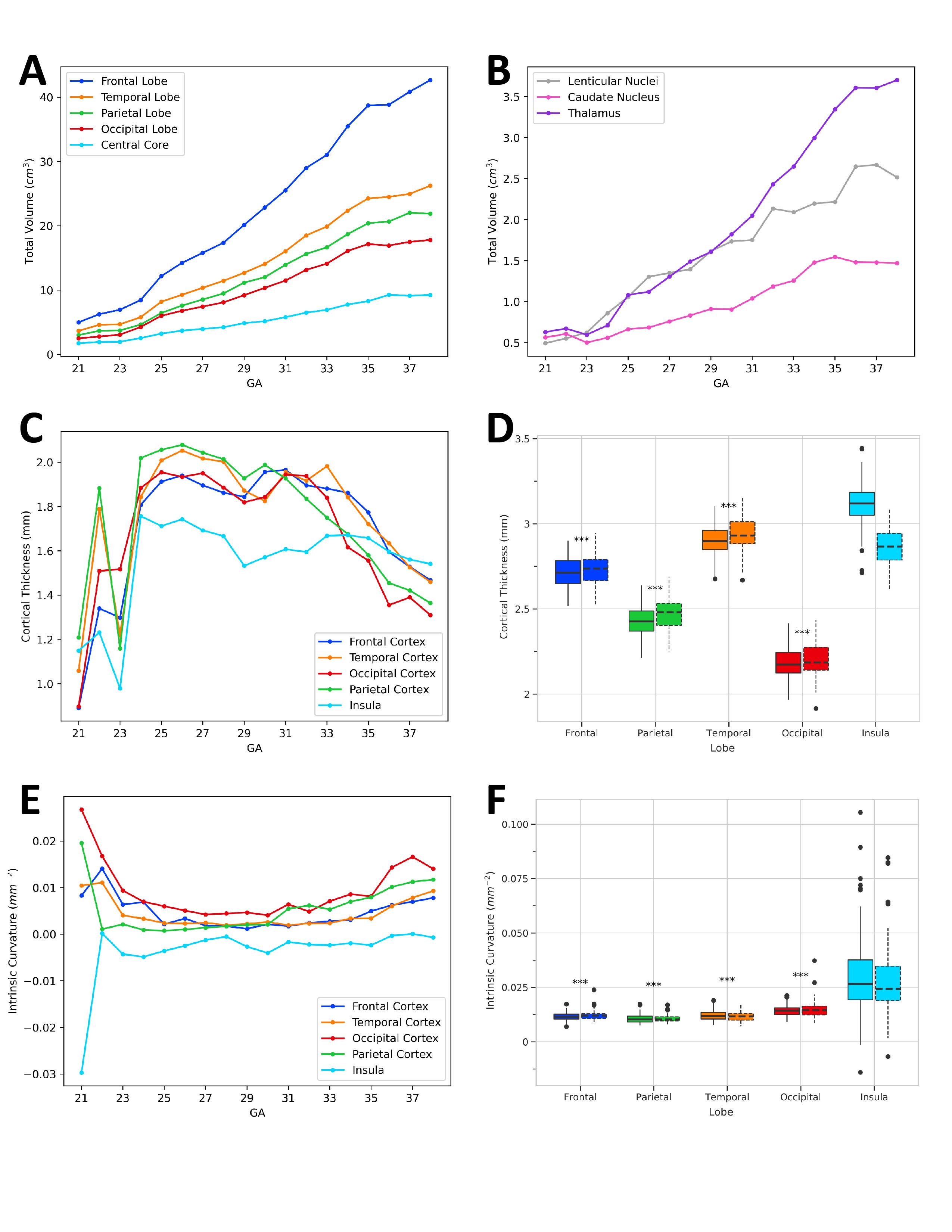
 **Figure S2:** **Volumetric analyses, cortical thickness, and intrinsic (Gaussian) curvature**. **A:** Volume of cerebral lobes (including white matter) and central core (insula + basal ganglia and deep white matter) by gestational age (GA). Growth curves are approximately logistic (see Supplementary Table 2). After logit transform, linear regression against side, GA, and lobe was significant with good overall fit (R^2^ = 0.953, F(6,173)=608.8, p<0.0001). Volume was significantly higher in the frontal (p < 0.0001), parietal (p < 0.0001), temporal (p < 0.0001), and occipital lobes (p < 0.0001). **B:** Volume of central core components by GA. The thalamus, as a diencephalic structure, was not considered part of the central core but is included for reference. **C:** Cortical thickness by lobe by gestational age during fetal development. Linear regression against side, GA, and lobe was significant but with poor overall fit (R^2^ = 0.124, F(6,173)=5.208, p<0.0001). Cortical thickness was only significantly greater in temporal cortex (p=0.008). **D:** Cortical thickness by lobe in adult cohort. Frontal, parietal, temporal, and occipital all had significantly thinner cortex than the insula (p < 0.001) **E:** Intrinsic (Gaussian) curvature by lobe by gestational age during fetal development. Linear regression against side, GA, and lobe was not significant (R^2^ = 0.044, F(6,173)=2.384, p=0.0308). Intrinsic curvature was significantly greater in parietal (p=0.008) and occipital (p<0.001) cortices. **F:** Intrinsic (Gaussian) curvature by lobe in adult cohort. Frontal, parietal, temporal, and occipital all had significantly less intrinsic curvature than the insula (p < 0.001, Bonferroni corrected) *** - p < 0.001 (Mann-Whitney Test – lobe vs. Insula, Bonferroni corrected)

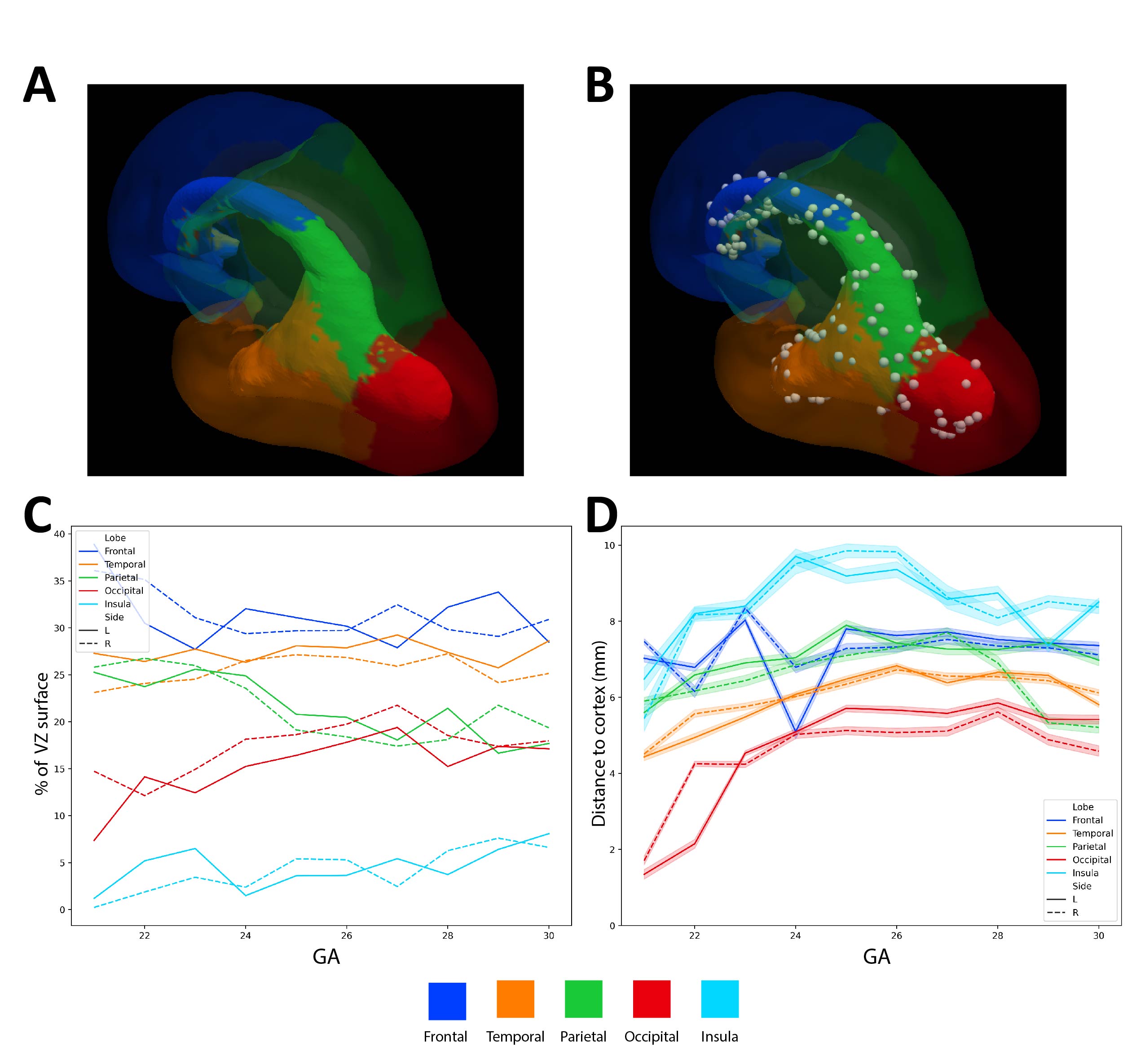
 **Figure S3:** **Segmentation of ventricular zone based off proximity to cortical surface. A:** Segmentation at GA 26 demonstrating large segments of VZ closest to frontal, temporal, parietal, and occipital cortical surfaces. Regions closest to insula are confined to the inferomedial area of frontal and parietal VZ and superomedial areas of temporal VZ. **B:** Random sampling of N=30 points for each VZ segmentation for projection analysis at same GA. **C:** Percentage of total VZ surface area closest to each cortical region. Linear regression against side, GA, and lobe was significant with good overall fit (R^2^ = 0.918, F(6,93)=186.4, p<0.0001). The proportion of VZ segmented to frontal, temporal, parietal, and occipital lobes was significantly higher compared to insula (p < 0.001). **D:** Distance from segmented VZ areas to respective lobe. Linear regression against side, GA, and lobe was significant but with poor overall fit (R^2^ = 0.172, F(6,93220)=3233, p<0.0001). Distance from VZ to cortical surface was significantly greater in frontal, temporal, parietal, and occipital VZ compared to insular VZ (p < 0.001)

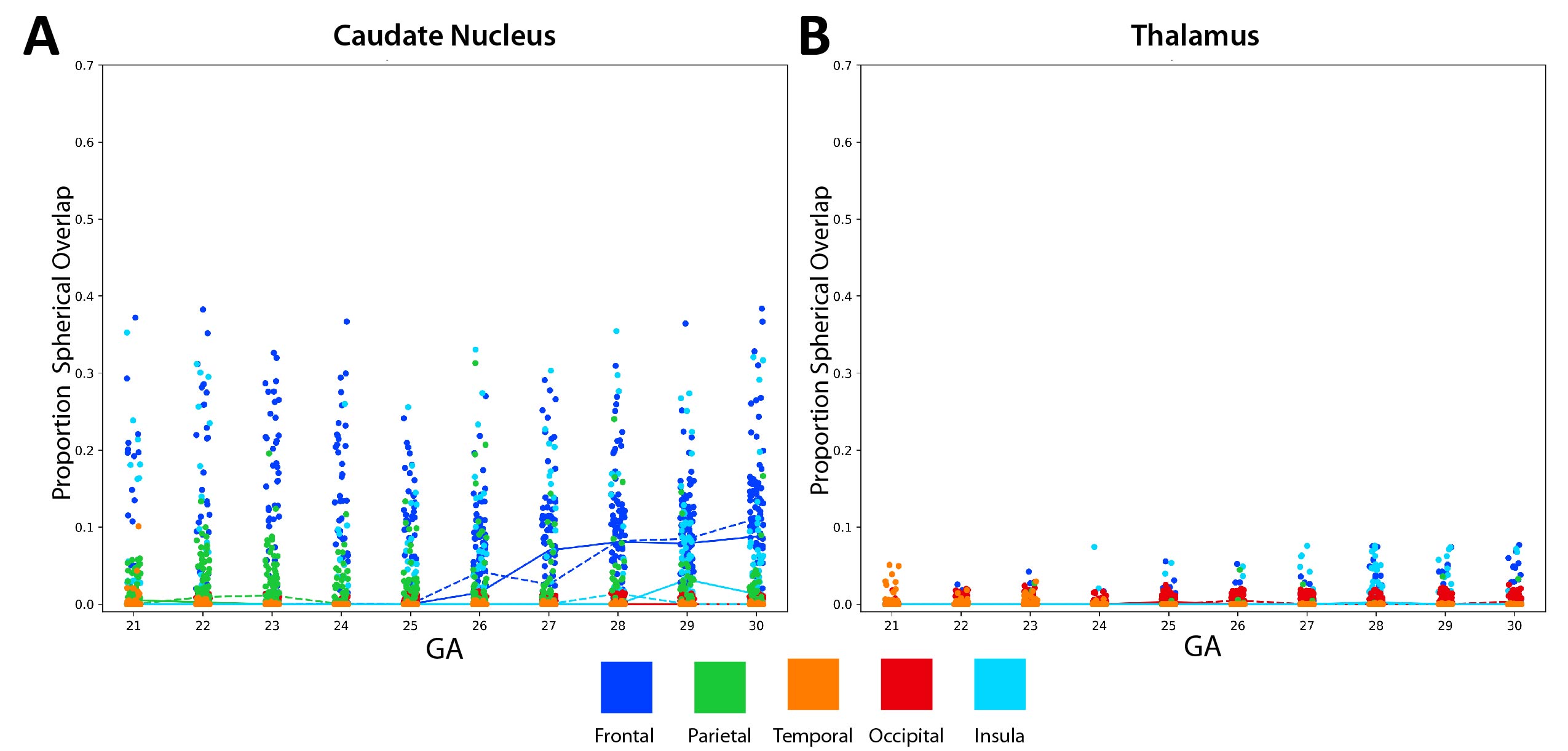
 **Figure S4: Results of spherical projection analysis for caudate nucleus and thalamus**. **A:** Caudate nucleus – the caudate nucleus overlaps (occludes) approximately a median of 0 to 10% of available radial paths from the VZ to the frontal lobe, and less than 5% of paths to the other lobes. Linear regression against side, GA, and lobe was significant but with poor overall fit (R^2^ = 0.215, F(6,4842)=222.6, p<0.0001). The proportion of spherical overlap was significantly higher in the frontal lobe than the insula (+4.0% [3.7-4.5], p < 0.01), and lower in the parietal (-1.0% [-1.4 - -0.6], p < 0.01), temporal (-2.2% [-2.6 - -1.8] p < 0.01), and occipital (-2.2% [-2.4 - -1.6] p < 0.01). The effect of GA was small but significant (0.1% per week, [0.1-0.2], p<0.01): **B**: Thalamus – the thalamus does not significantly occlude any path from the ventricular zone to the cortical surface, consistent with its diencephalic origin. Linear regression against side, GA, and lobe was significant but with poor overall fit (R^2^ = 0.038, F(6,4842)=33.29, p<0.0001). The proportion of spherical overlap was significantly lower in the parietal (-0.2% [-0.3 - -0.2], p < 0.01) and temporal (-0.2% [-0.3 - -0.1], p < 0.01) lobes than the insula, but not significantly different between the frontal/occipital lobes and the insula. GA had a negligible effect on overlap

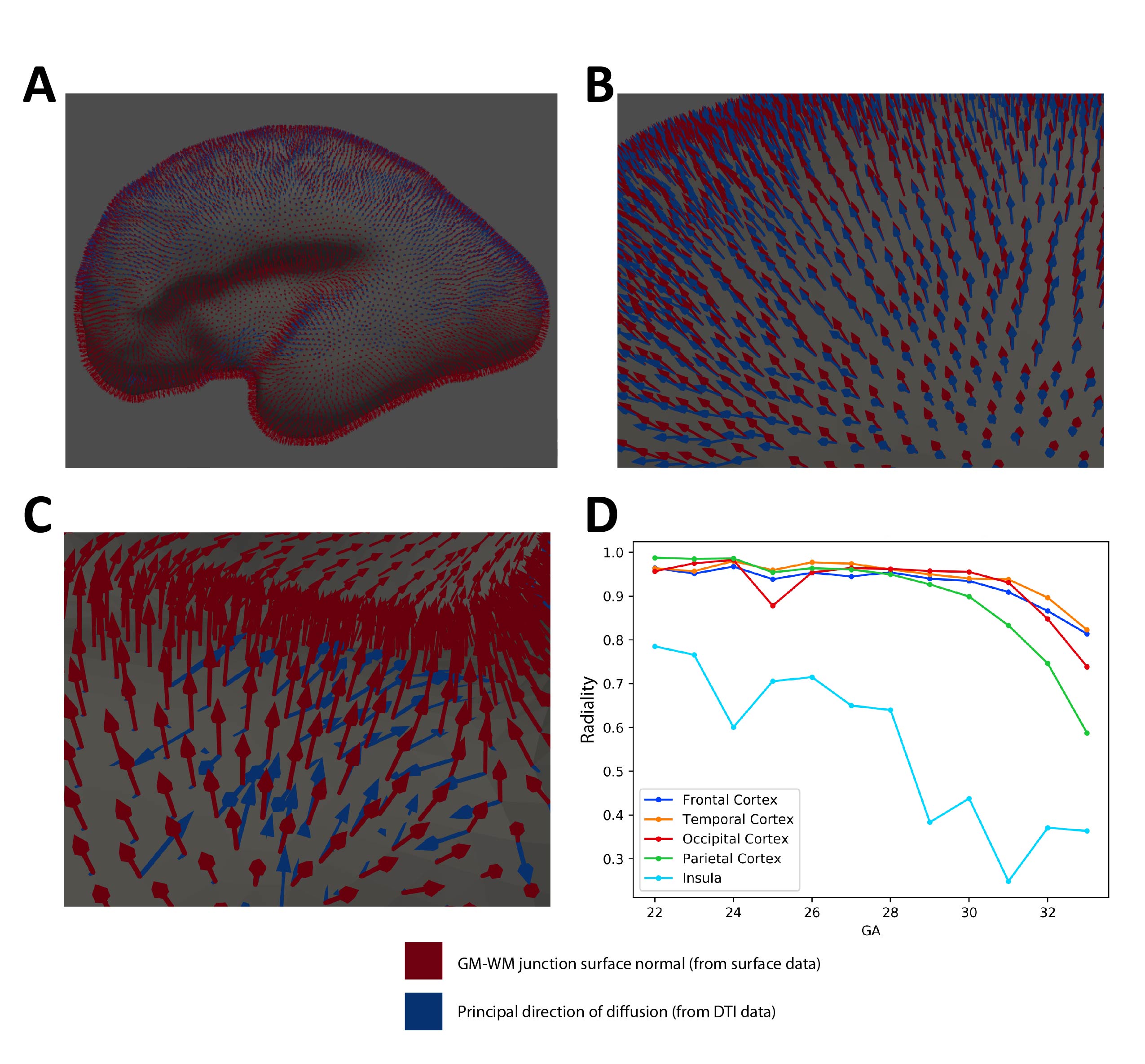

**Figure S5: Radiality analysis**. Radiality, as defined by Eaton-Rosen 2017, measures the degree to which migrating glial cells or white matter is perpendicular to the cortical surface. Their analysis demonstrates that radiality declines in the frontal, parietal, temporal, and occipital cortices in a GA-dependent fashion starting at approximately 27 weeks. Notably, the insula was not included. Radiality is defined as the normalized dot product of the surface normal ***r*** and the principal eigenvector of the diffusion tensor (principal direction of diffusion), ***v***, e.g.:

$$\frac{\left| \boldsymbol{r \cdot v} \right|}{\left\| \boldsymbol{r} \right\|\left\| \boldsymbol{v} \right\|}$$

Surface normal were calculated for the gray matter – white matter junction surface. **A:** Example of surface normals (red) and the principal direction of diffusion (blue) for the left hemisphere at GA 24. **B:** Frontal lobe surface. Note how the surface normal and principal direction of diffusion are well-aligned, resulting in a high radiality. **C:** Insular surface. Surface normal and principal direction of diffusion are not well-aligned (almost perpendicular), resulting in low radiality. **D:** Extended version of Figure 4E, demonstrating the linear decline in ROI after GA 27-28, consistent with the findings of Eaton-Rosen 2017.

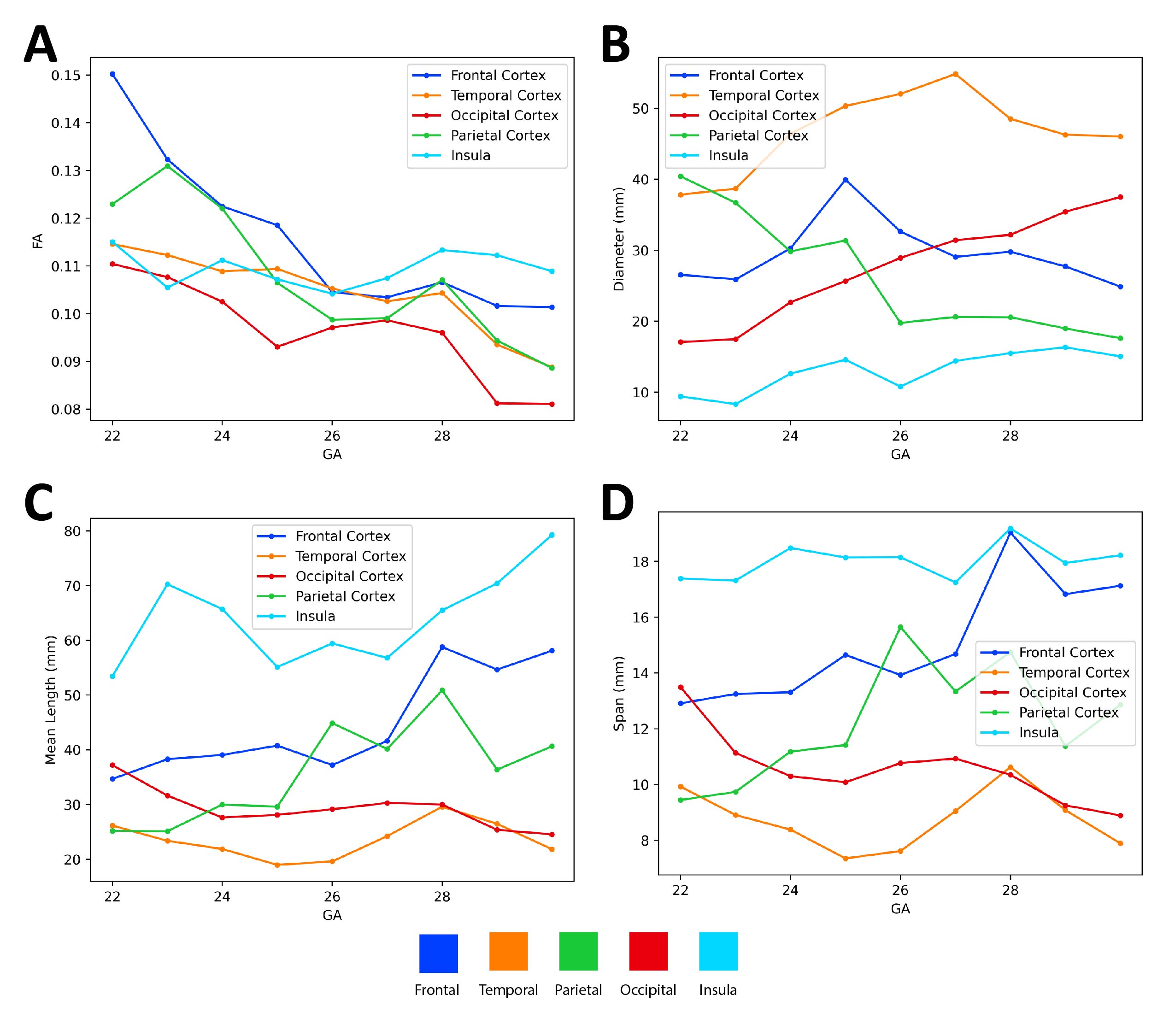
 **Figure S6**: **Shape analysis of radial glial fascicles from diffusion tractography.** Using the entire VZ as an end and cortical surface as region of interest (by lobe), we identified putative streams of developing cells from the ventricular zone to cortical surface. We used shape analysis of diffusion tracts (described by Yeh 2020) to examine the fascicles to each lobe **A:** Mean fractional anisotropy (FA) per set of fascicles from the VZ to each lobe by GA. Linear regression against side, GA, and lobe was significant with moderate overall fit (R^2^ = 0.594, F(6,83)=22.69, p<0.0001). Mean FA was significantly lower for fascicles going to the occipital lobe (p<0.001), but not between those going to the frontal/temporal/parietal lobes and the fascicles to the insula. **B:** Mean fascicle diameter per set of fascicles from the VZ to each lobe by GA. Linear regression against side, GA, and lobe was significant with good overall fit (R^2^ = 0.740, F(6,83)=43.13, p<0.0001). Diameter was significantly larger for fascicles going to all other lobes (frontal, temporal, parietal, occipital) lobe than to the insula (p<0.001). **C:** Mean length (curvilinear distance along tract) per set of fascicles from the VZ to each lobe by GA. Linear regression against side, GA, and lobe was significant with good overall fit (R^2^ = 0.758, F(6,83)=47.58, p<0.0001). Mean length was significantly smaller for fascicles going to all other lobes (frontal, temporal, parietal, occipital) lobe than to the insula (p<0.001). **D:** Mean span (Euclidean distance from tract terminus to terminus) per set of fascicles from the VZ to each lobe by GA. Linear regression against side, GA, and lobe was significant with good overall fit (R^2^ = 0.763, F(6,83)=48.69, p<0.0001). Mean span was significantly smaller for fascicles going to all other lobes (frontal, temporal, parietal, occipital) lobe than to the insula (p<0.001).

**Table S1**: ID and basic information for adults from HCP 900 release used for adult analysis. N=107, 54.2% Female, Median Age = 28

| Subject | Release | Acquisition | Gender | Age |
| --- | --- | --- | --- | --- |
| 100206 | S900 | Q11 | M | 26-30 |
| 100610 | S900 | Q08 | M | 26-30 |
| 102513 | S900 | Q10 | M | 26-30 |
| 104416 | S900 | Q09 | F | 31-35 |
| 105620 | S900 | Q08 | F | 31-35 |
| 107018 | S900 | Q09 | F | 26-30 |
| 107220 | S900 | Q11 | F | 26-30 |
| 107725 | S900 | Q08 | F | 31-35 |
| 108222 | S900 | Q07 | M | 31-35 |
| 109830 | S900 | Q08 | F | 31-35 |
| 110007 | S900 | Q11 | F | 31-35 |
| 110613 | S900 | Q11 | M | 26-30 |
| 111009 | S900 | Q03 | F | 26-30 |
| 112112 | S900 | Q11 | M | 26-30 |
| 112314 | S900 | Q11 | F | 26-30 |
| 112516 | S900 | Q08 | F | 31-35 |
| 112920 | S900 | Q10 | M | 31-35 |
| 114217 | S900 | Q08 | F | 26-30 |
| 114318 | S900 | Q06 | F | 22-25 |
| 114621 | S900 | Q09 | M | 26-30 |
| 114823 | S900 | Q07 | F | 31-35 |
| 115017 | S900 | Q06 | F | 31-35 |
| 115219 | S900 | Q09 | M | 31-35 |
| 115825 | S900 | Q10 | M | 22-25 |
| 116221 | S900 | Q08 | M | 22-25 |
| 116726 | S900 | Q08 | M | 26-30 |
| 117930 | S900 | Q10 | F | 31-35 |
| 118023 | S900 | Q07 | F | 26-30 |
| 118124 | S900 | Q07 | F | 31-35 |
| 118225 | S900 | Q08 | M | 26-30 |
| 119126 | S900 | Q11 | F | 22-25 |
| 119732 | S900 | Q10 | F | 31-35 |
| 120717 | S900 | Q09 | F | 31-35 |
| 121416 | S900 | Q08 | M | 26-30 |
| 121820 | S900 | Q07 | F | 31-35 |
| 121921 | S900 | Q07 | M | 31-35 |
| 122822 | S900 | Q07 | F | 31-35 |
| 123521 | S900 | Q10 | M | 22-25 |
| 123824 | S900 | Q11 | M | 22-25 |
| 124624 | S900 | Q09 | F | 26-30 |
| 127327 | S900 | Q08 | M | 22-25 |
| 128026 | S900 | Q10 | F | 26-30 |
| 128935 | S900 | Q09 | M | 31-35 |
| 129129 | S900 | Q07 | F | 31-35 |
| 129331 | S900 | Q09 | F | 26-30 |
| 129634 | S900 | Q11 | M | 22-25 |
| 129937 | S900 | Q11 | F | 26-30 |
| 130417 | S900 | Q07 | M | 26-30 |
| 130619 | S900 | Q10 | F | 26-30 |
| 130821 | S900 | Q10 | F | 22-25 |
| 131419 | S900 | Q09 | F | 26-30 |
| 131823 | S900 | Q11 | M | 26-30 |
| 132017 | S900 | Q08 | F | 22-25 |
| 134021 | S900 | Q10 | F | 31-35 |
| 134223 | S900 | Q11 | F | 26-30 |
| 134425 | S900 | Q07 | F | 26-30 |
| 134728 | S900 | Q08 | M | 26-30 |
| 134829 | S900 | Q11 | F | 31-35 |
| 135730 | S900 | Q09 | M | 22-25 |
| 136732 | S900 | Q07 | F | 31-35 |
| 137229 | S900 | Q10 | M | 22-25 |
| 138837 | S900 | Q11 | M | 22-25 |
| 139839 | S900 | Q07 | M | 26-30 |
| 140319 | S900 | Q08 | F | 36+ |
| 141119 | S900 | Q09 | M | 26-30 |
| 143426 | S900 | Q11 | F | 26-30 |
| 144125 | S900 | Q12 | F | 31-35 |
| 144731 | S900 | Q10 | F | 26-30 |
| 145127 | S900 | Q08 | M | 22-25 |
| 146533 | S900 | Q12 | M | 26-30 |
| 146634 | S900 | Q04 | F | 26-30 |
| 146937 | S900 | Q08 | F | 31-35 |
| 148133 | S900 | Q10 | F | 26-30 |
| 148436 | S900 | Q08 | M | 26-30 |
| 149236 | S900 | Q08 | F | 22-25 |
| 149842 | S900 | Q10 | F | 31-35 |
| 150019 | S900 | Q12 | M | 26-30 |
| 150928 | S900 | Q11 | M | 22-25 |
| 151425 | S900 | Q09 | F | 26-30 |
| 151829 | S900 | Q10 | F | 31-35 |
| 153227 | S900 | Q12 | F | 31-35 |
| 153631 | S900 | Q10 | F | 22-25 |
| 154229 | S900 | Q10 | F | 22-25 |
| 154532 | S900 | Q09 | M | 26-30 |
| 155938 | S900 | Q08 | M | 26-30 |
| 156031 | S900 | Q11 | F | 26-30 |
| 156435 | S900 | Q11 | M | 22-25 |
| 156536 | S900 | Q12 | M | 22-25 |
| 157942 | S900 | Q07 | F | 36+ |
| 158338 | S900 | Q08 | M | 22-25 |
| 158843 | S900 | Q08 | M | 26-30 |
| 159744 | S900 | Q11 | M | 31-35 |
| 159845 | S900 | Q11 | F | 31-35 |
| 159946 | S900 | Q07 | M | 26-30 |
| 160729 | S900 | Q08 | F | 22-25 |
| 160931 | S900 | Q08 | F | 31-35 |
| 162935 | S900 | Q09 | M | 22-25 |
| 164636 | S900 | Q09 | M | 22-25 |
| 165638 | S900 | Q07 | M | 26-30 |
| 166640 | S900 | Q10 | M | 22-25 |
| 167238 | S900 | Q08 | F | 31-35 |
| 168038 | S900 | Q08 | M | 26-30 |
| 168240 | S900 | Q08 | M | 26-30 |
| 168745 | S900 | Q11 | F | 22-25 |
| 169040 | S900 | Q08 | M | 22-25 |
| 169747 | S900 | Q10 | F | 26-30 |
| 169949 | S900 | Q11 | M | 26-30 |

**Table S2**: Best model fits and parameters for volume and surface metrics by gestational age.

| **Metric** |  | **Frontal** | **Temporal** | **Occipital** | **Parietal** | **Insula/Central Core** |
| --- | --- | --- | --- | --- | --- | --- |
| **Volume (cm^3^)** | **Model Type** | Logistic | Logistic | Logistic | Logistic | Logistic |
|  | **AIC** | -4.31 | -14.38 | -21.39 | -27.56 | -84.22 |
|  | **Parameters** | α = 0.05; K = 51.9;  N_0_ = 0.22 | α = 0.04; K = 32.18;  N_0_ = 0.22 | α = 0.02; K = 20.94;  N_0_ = 0.23 | α = 0.03; K =26.76;  N_0_= 0.23 | α = 0.002; K = 4.08;  N_0_ = 0.23 |
| **Surface Area (cm^2^)** | **Model Type** | Exponential | Exponential | Exponential | Exponential | Linear |
|  | **AIC** | 41.33 | 6.77 | 39.00 | 41.33 | -43.88 |
|  | **Parameters** | α = 0.107; N_0_ = 1.811 | α = 0.108; N_0_ = 1.013 | α = 0.102; N_0_ = 1 | α = 0.114; N_0_ = 1 | m = 0.042; N_0_ = -8.11 |
| **Cortical Thickness (mm)** | **Model Type** | Linear | Linear | Linear | Linear | Linear |
|  | **AIC** | -42.57 | -42.57 | -40.60 | -40.51 | -56.41 |
|  | **Parameters** | m = 0.016; N_0_ = 1.229 | m = 0.004; N_0_ = 1.633 | m = -0.004; N_0_ = 1.799 | m = -0.013; N_0_ = 2.116 | m = 0.018; N_0_ = 1.034 |
| **Sulcal depth range (cm)** | **Model Type** | Linear | Linear | Linear | Linear | Linear |
|  | **AIC** | -170.82 | -176.32 | -155.22 | -157.75 | -203.58 |
|  | **Parameters** | m = 0.005; N_0_ = 0.009 | m = 0.008; N_0_ = -0.052 | m = 0.006; N_0_ = -0.013 | m = 0.009; N_0_ = -0.086 | m = 0.005; N_0_ = -0.042 |
| **Mean curvature (mm^-1^)** | **Model Type** | Linear | Linear | Linear | Linear | Linear |
|  | **AIC** | -152.56 | -170.59 | -134.74 | -174.26 | -156.94 |
|  | **Parameters** | m = -0.004; N_0_ = 0.162 | m = -0.002; N_0_= 0.114 | m = -0.002; N_0_ = 0.109 | m = -0.002; N_0_ = 0.089 | m = 0.000; N_0_ = -0.043 |
| **Gaussian curvature (mm^-2^)** | **Model Type** | Linear | Linear | Linear | Linear | Linear |
|  | **AIC** | -203.11 | -205.03 | -180.89 | -189.18 | -181.92 |
|  | **Parameters** | m = -0.000; N_0_ = 0.009 | m = -0.000; N_0_ = 0.005 | m = -0.000; N_0_ = 0.012 | m = 0.000; N_0_ = -0.004 | m = 0.001; N_0_ = -0.021 |
| **Fractal Dimension (unitless)** | **Model Type** | Linear | Linear | Linear | Linear | Linear |
|  | **AIC** | -85.302232 | -91.479803 | -97.634039 | -87.502684 | -96.532984 |
|  | **Parameters** | m = 0.030; N_0_ = 1.225 | m = 1.397; N_0_ = 0.014 | m = 0.026; N_0_ = 1.309 | m = 0.030; N_0_ = 1.18 | m = 0.028; N_0_ = 0.975 |

**Formulas:**

*GA –* Gestational Age (range 21 – 38 weeks)

Linear:

$$Metric=m*GA+N_{0}$$

Exponential:

$$Metric=N_{0}* e^{\alpha*GA}$$

Logistic:

$$Metric=\frac{N_{0}*K}{N_{0}+\left( K- N_{0} \right)* e^{-\alpha*GA}}$$

**Table S3**: Linear fits for DTI Radiality from GA22-33. Note similar values to Eaton-Rosen 2017, but smaller intercept and more negative slope for insula than other lobes

| **Lobe** | **DTI Radiality** |  |
| --- | --- | --- |
|  | **Intercept ± SE (27 weeks)** | **Slope ± SE (per week)** |
| **Frontal** | 0.927 ± 0.008 | -0.011 ± 0.002 |
| **Temporal** | 0.94 ± 0.009 | -0.011 ± 0.002 |
| **Occipital** | 0.932 ± 0.01 | -0.011 ± 0.003 |
| **Parietal** | 0.918 ± 0.013 | -0.026 ± 0.004 |
| **Insula** | 0.607 ± 0.022 | -0.032 ± 0.006 |
